# Alveolar Repair Following Lipopolysaccharide-induced Injury Requires Cell-Extracellular Matrix Interactions

**DOI:** 10.1101/2022.08.05.502988

**Authors:** Jennifer M.S. Sucre, Fabian Bock, Nicholas M. Negretti, John T. Benjamin, Peter M. Gulleman, Xinyu Dong, Kimberly T. Ferguson, Christopher S. Jetter, Wei Han, Yang Liu, Seunghyi Kook, Jason J. Gokey, Susan H. Guttentag, Jonathan A. Kropski, Timothy S. Blackwell, Roy Zent, Erin J. Plosa

**Author notes:** Correspondence: Erin Plosa, MD, 2215 B Garland Ave, 1125 Light Hall/ MRB IV, Nashville, TN 37232-0656.

## Abstract

During alveolar repair, alveolar type 2 (AT2) epithelial cell progenitors rapidly proliferate and differentiate into flat type 1 alveolar epithelial cells. Failure of normal alveolar repair mechanisms can lead to loss of alveolar structure (emphysema) or development of fibrosis, depending on the type and severity of injury. To test if β1-containing integrins are required during repair following acute injury, we administered *E. coli* lipopolysaccharide (LPS) by intratracheal injection to mice with a post-developmental deletion of β1 integrin in AT2 cells. While control mice recovered from LPS injury without structural abnormalities, β1-deficient mice had more severe inflammation and developed emphysema. In addition, recovering alveoli were repopulated with an abundance of rounded epithelial cells co-expressing type 2, type 1, and mixed intermediate cell state markers, with few mature type 1 cells. β1-deficient AT2 cells showed persistently increased proliferation after injury, which was blocked by inhibiting NF-κB activation in these cells. Lineage tracing experiments revealed that β1-deficient AT2 cells failed to differentiate into mature type 1 alveolar epithelial cells. Together, these findings demonstrate that functional alveolar repair after injury with terminal alveolar epithelial differentiation requires β1-containing integrins.

## Introduction

During lung development, primordial epithelial cells proliferate, migrate, and change phenotypic identity with precise timing, anchored by signals from the basement membrane (BM), a specialized extracellular matrix (ECM) structure, that is precisely remodeled at specific developmental check points (1). Once lung development is complete, targeted replacement of epithelial cells and slow turnover of the alveolar basement membrane protects the alveolar structure during homeostasis in the mature lung. However, epithelial cells lose capacity for efficient proliferation and differentiation as the BM ages (2), increasing susceptibility to chronic lung diseases over time. In contrast to temporally precise development and the relatively quiescent adult lung, repair of acute lung injury must occur rapidly in order to restore a gas-exchanging epithelium to maintain survival. The immediacy of repair results in epithelial proliferation and differentiation that occurs in bulk. For a wild-type mouse, the mild lung injury induced by a single dose of intratracheal lipopolysaccharide (LPS) is easily recoverable in days, overcoming inflammation-derived proteolytic damage to the basement membrane that occurs within hours (3, 4). LPS, as a model of acute inflammatory lung injury, tests the regenerative potential of the alveolus, exposing impairments in epithelial-ECM interactions that could exacerbate lung injury or predispose to accelerated aging.

Integrins are heterodimeric transmembrane protein receptors composed of an α and β subunit that bind ECM ligands. Integrins provide physical connections between cells and the ECM, and they propagate signaling to and from the surrounding matrix (5–7). Of the 24 integrin heterodimers, 12 contain the β1 subunit, and many of the 12 β1-containing integrins are present in epithelial tissues. Integrin function is dependent upon developmental and microenvironmental context, a concept consistent with our previous work. We previously reported that epithelial β1 integrins are required for normal lung development and in their absence airway branching and alveolarization are impaired, associated with incomplete epithelial differentiation in the later stages of lung development (8). However, in the homeostatic post-developmental adult alveolus, β1 integrins are completely dispensable for alveolar epithelial differentiation (9). Function of the alveolar stem cell niche during repair depends on the rapid proliferation of distal epithelial progenitors, followed by en masse differentiation. The degree of differentiation in this context is profound, and any inefficiencies in this process will create structural vulnerability and abnormal repair. Whether or not β1-containing integrins are required for the epithelial repair cycle in the acutely injured adult alveolus is unknown.

The adult alveolar BM is primarily composed of collagen IV and laminin and facilitates gas exchange between the airspace and the capillary network. The vast majority of the BM is covered with very thin and outspread type 1 alveolar epithelial cells (AT1 cells), interspersed with single cuboidal type 2 alveolar epithelial cells (AT2 cells). Each cell type has essential functions: AT2 cells secrete surfactant to reduce alveolar surface tension, and function in an immunoregulatory role, while the thin AT1 cells allow gas exchange, maintain water homeostasis, and produce BM components (10). Following alveolar injury in both human lung disease and murine models, multiple types of epithelial progenitors restore epithelial coverage of the BM (11, 12). The most proximate and immediate epithelial progenitors for alveolar repair are AT2 cells themselves, a subset of which proliferate upon alveolar injury (13–15). Over-proliferation of AT2 cells is restrained by various proteins including the tight junction protein Claudin-18 (16). After proliferation, the AT2 progenitors attain an AT2-to-AT1 transitional cell state with characteristic expression of genes such as *Krt8, Hbegf,* or *Areg* (11, 17, 18), on their way to becoming replacement AT1 cells. Less is understood about the molecular mechanisms that govern the final point of transition in alveolar repair, the acquisition of the AT1 transcriptional phenotype and the full extension of the AT1 outspread cell shape.

Here, we compared distal lung epithelium restoration after acute lung injury induced by intratracheal LPS administration in normal adult mice to those carrying a β1-integrin inactivation targeted to SP-C–expressing AT2 cells. From this modest acute lung injury, β1-deficient mice exhibited increased inflammation and had exacerbated lung injury, followed by emphysema. Impaired alveolar repair mechanisms in injured β1-deficient mice resulted in excessive AT2 proliferation, progressive accumulation of AT2/AT1 intermediate cells, and large-scale failure of AT1 differentiation. In these studies, the degree of cell-shape block in β1-deficient mice demonstrates that AT2-to-AT1 differentiation is tightly integrated with cell shape change. Further, we show that epithelial differentiation during repair bears a particular requirement for β1 integrin-mediated cell-ECM interactions, more than AT2-to-AT1 differentiation in other contexts. Unlike the assumption that AT2-to-AT1 epithelial differentiation processes employ similar basic mechanistic pathways no matter the context, our data suggest a different paradigm for adult injury repair wherein the rapidity with which differentiation must occur and/or the condition of the ECM leads to distinct differentiation mechanisms.

## Results

### β1 integrin deletion in AT2 cells increases susceptibility to injury and abnormal repair

To determine the role of epithelial β1 integrin in alveolar repair after injury, we generated mice with a β1-integrin deletion in AT2 cells in the adult lung utilizing the SP-C–rtTA; TetO-Cre doxycycline-inducible system (9, 19) and floxed β1-integrin (β1^f/f^), hereafter referred to as β1^AT2-KO^ mice. We previously reported the efficient deletion of β1 integrin in AT2 cells and sustained β1 integrin expression in β1^f/f^ littermate control mice (then termed β1^rtTA^ mice) and confirmed normal histology in SP-C rtTA; TetO-Cre; β1^f/f^ mice not receiving doxycycline at three months of age (9, 19). While doxycycline toxicity has been reported with embryonic administration (20), we observed no overt histologic or biological toxicity from the rtTA system or from doxycycline itself in this post-developmental model (9). After one month off doxycycline to minimize side effects from possible doxycycline tissue storage, we challenged β1^AT2-KO^ and littermate β1^f/f^ controls with a single intratracheal (IT) dose of lipopolysaccharide (LPS) at three months of age (**Figure 1A**). β1^AT2-KO^ mice exhibited decreased survival compared to β1^f/f^ control mice (44% survival at 21 days for β1^AT2-KO^ versus 93% for β1^f/f^, n=15 β1^AT2-KO^ and 14 β1^f/f^ mice, *p*=0.0051, **Supp. Figure 1A**), with 100% survival for both PBS-treated β1^AT2-KO^ and β1^f/f^ mice (data not shown). Three days post-injury, histological examination revealed an expected increase in inflammatory cells in both LPS-treated β1^AT2-KO^ and control β1^f/f^ lungs and markedly increased edema in β1^AT2-KO^ lungs (Figure 1A). Whereas LPS-treated β1^f/f^ lungs returned to a normal histological appearance by day 7, edema and inflammation remained prominent in β1^AT2-KO^ lungs, consistent with an exacerbation of acute lung injury. Failed alveolar repair was evident at 21 days in β1^AT2-KO^ mice, whose lungs exhibited a marked increase in airspace size and septal destruction. Emphysema was quantified by increased mean linear intercept (28.5±0.9 μm in β1^f/f^lungs vs. 40.2±2.8 μm in β1^AT2-KO^ lungs) (**Figure 1B**). To evaluate alveolar barrier function, we measured bronchoalveolar lavage (BAL) fluid protein levels at all time points (**Figure 1C**). β1^AT2-KO^ BAL fluid contained increased protein in the unchallenged state, as well as at three and seven days post-LPS compared to control β1^f/f^ BAL fluid. By 21 days post-injury, protein levels were similar in both mouse strains, indicating restoration of alveolar barrier function in β1^AT2-KO^ lungs.

**Figure 1.**
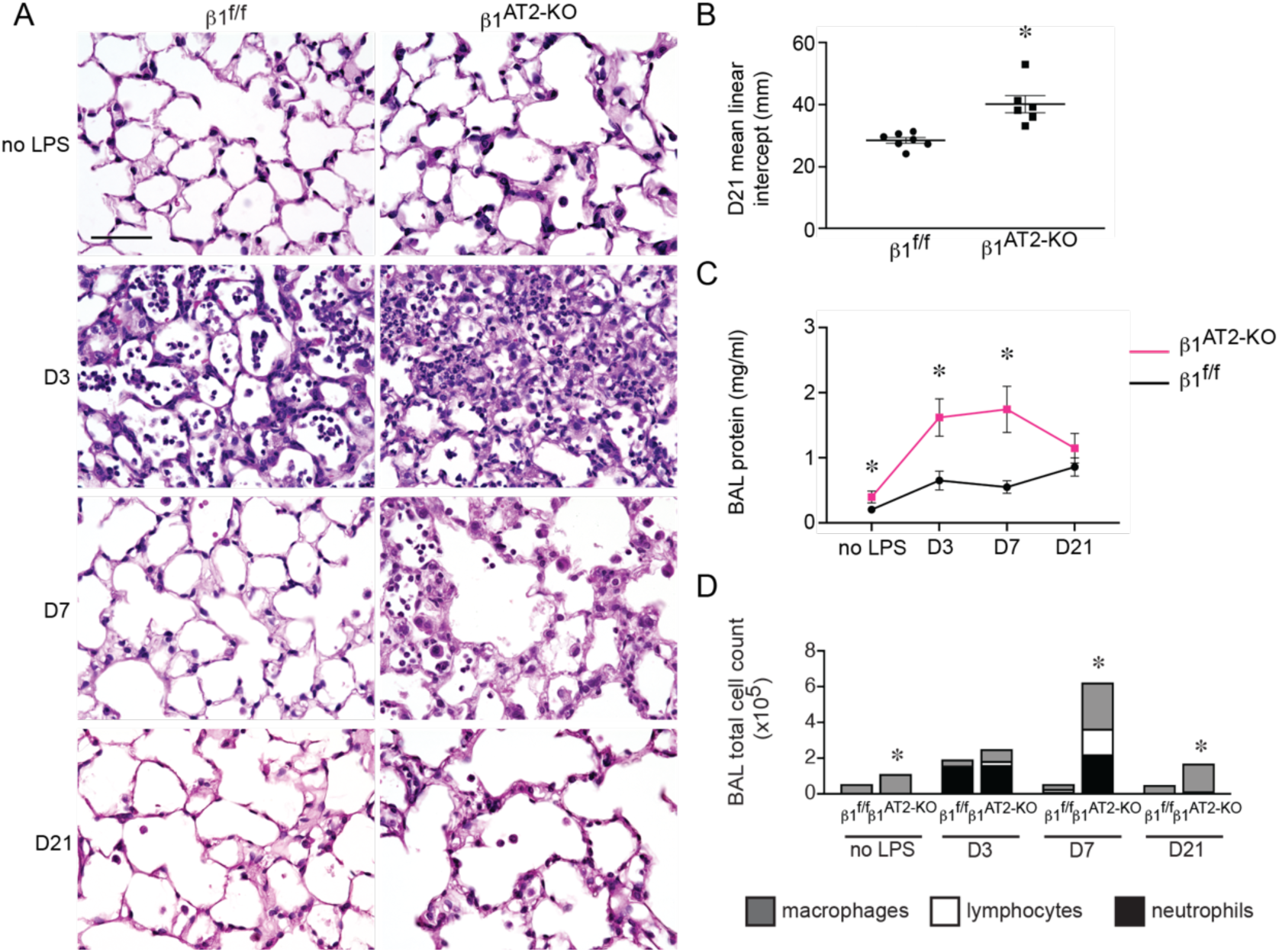
β1 integrin deletion in AT2s results in increased inflammation, abnormal repair, and decreased survival after LPS-induced injury. (**A**) Representative images of lung histology demonstrate increased edema at 3 and 7 days post-injury in β1^AT2-KO^ lungs compared to β1^f/f^ lungs, as well as persistent inflammation and emphysematous remodeling in β1^AT2-KO^ lungs by 21 days post-LPS. -LPS refers to uninjured mice; D3, D7, and D21 refer to 3, 7, and 21 days post-LPS injury, respectively. (**B**) Mean linear intercept quantified emphysematous alveolar remodeling at 21 days post-LPS, 28.5±0.9 μm in β1^f/f^ lungs vs. 40.2±2.8 μm in β1^AT2-KO^ lungs (n=6-7 mice/ group, *p*=0.0014 by two-tailed t-test). (**C**) Bicinchoninic Acid protein assay quantified increased bronchoalveolar lavage (BAL) fluid protein in uninjured β1^AT2-KO^ lungs and at 3 and 7 days post-LPS injury in β1^AT2-KO^ lungs compared to β1^f/f^ lungs at the same time points (n=6-14 mice/ group, two-tailed t-test comparing genotypes at each time point, *p*=0.0485 for uninjured mice; *p*=0.0036 at D3; *p*=0.005 at D7; *p*=0.2628 at D21). (**D**) BAL cell counts are significantly increased in β1^AT2-KO^ lungs compared to β1^f/f^ littermates in uninjured mice and at 7 and 21 days post-LPS. Peak inflammation is present at 7 days in β1^AT2-KO^ lungs, 55,663±3306 cells/ ml in β1^f/f^ BALF versus 624,000±118,753 cells/ml in β1^AT2-KO^ BAL (n=6-26 mice/group, two-tailed t-test comparing genotypes at each time point, *p*=0.0002 for uninjured mice; *p*=0.0730 at D3; *p*=0.0007 at D7; *p*<0.0001 at D21). Total numbers of BALF macrophages are significantly increased in uninjured β1^AT2-KO^ mice and at day 7 and day 21, lymphocytes and neutrophils are significantly increased in β1^AT2-KO^ BAL at day 7 only. Scale bar=50 μm for (A). * *p*< 0.05.

We collected BAL fluid to characterize the inflammatory response to LPS and found no significant differences in immune/ inflammatory cells between β1^AT2-KO^ and β1^f/f^ lungs at three days, during the expected peak of inflammation following LPS (21) (**Figure 1D**). However, in β1^AT2-KO^ mice, there was a delayed inflammatory peak at seven days post-injury characterized by persistently increased neutrophils in BAL fluid from β1^AT2-KO^ lungs (**Supp. Figure 1B**). Although the BAL cell count decreased towards baseline at 21 days in β1^AT2-KO^ lungs, increased numbers of macrophages remained. These findings were confirmed by quantification of immunostaining for the macrophage marker CD68 (**Supp. Figure 1C-D**). Since chronic inflammation predisposes to emphysema, we tested if the exaggerated acute inflammatory response contributed to airspace expansion in LPS-treated β1^AT2-KO^ lungs. We delayed doxycycline administration for five days after IT LPS dosing, thereby inducing β1 integrin deletion after acute inflammation. Lungs were harvested 21 days after LPS treatment (and 16 days after the initiation of doxycycline). On histological examination, β1^AT2-KO^ lungs retained emphysematous structural deficits, quantified by an increased mean linear intercept (**Supp. Figure 1E-F**).

We next investigated whether LPS-treated β1^AT2-KO^ mice exhibited fibrosis, an additional manifestation of abnormal alveolar repair apart from emphysema. By Masson’s trichrome stain, β1^AT2-KO^ and β1^f/f^ lungs did not exhibit gross differences in fibrillar collagen deposition (**Supp. Figure 2A**). The alveolar basement membrane also appeared similarly composed between genotypes, with comparable pan-laminin localization in the alveolar septum 21 days after LPS treatment (**Supp. Figure 2B**). As loss of one integrin may result in the upregulation of alternative integrin subunits, we examined LPS-treated β1^AT2-KO^ and β1^f/f^ lungs for differences in RGD-binding integrins. In this subgroup of integrins, heterodimers comprised of varied β subunits, bind ECM proteins containing the highly conserved Arg-Gly-Asp (RGD) motif (22). We first confirmed continued absence of β1 at 21 days after LPS in β1^AT2-KO^ AT2 cells (Supp. Figure 2B, arrows). Integrin α5, which partners only with β1 to form a fibronectin receptor, was also decreased throughout injured β1^AT2-KO^ lungs compared to β1^f/f^ lungs (**Supp. Figure 3A**). In contrast to α5β1, the αv RGD-binding integrins were increased (**Supp. Figure 3B-E**), particularly the β5 and β8 subunits, suggesting compensation by αvβ5 and αvβ8 without overt fibrosis in the injured β1^AT2-KO^ lung. Taken together, these findings indicate that loss of β1-containing epithelial integrins drive increased LPS-induced inflammation and lung injury, followed by emphysema and abnormal alveolar repair.

### β1-deficient AT2 cells increase in number post-LPS injury

To determine how β1-deficiency in AT2 cells alters alveolar repair, we performed an in-depth histological and transcriptomic examination of alveolar epithelial cells in β1^AT2-KO^ and β1^f/f^ lungs. Lung sections immunostained for the AT2 marker pro-SP-C and AT1 marker T1α demonstrate accumulation of pro-SP-C+ AT2s in β1^AT2-KO^ lungs by day seven following LPS injury (**Supp. Figure 4A**). By day 21, β1^AT2-KO^ lungs contained increased numbers of pro-SP-C+ AT2 cells throughout the lung parenchyma (**Figure 2A-B**, quantified in **2C**). Moreover, severely injured areas exhibited alveolar septa lined by abundant pro-SP-C+ AT2s (Supp. Figure 4A). The increase in AT2 number was accompanied by decreased expression of the AT1 markers T1α and HOPX in β1^AT2-KO^ lungs (**Figure 2A-B**, HOPX quantified in **2D**), suggesting loss of AT1s and large expansion of the AT2 population. We captured the increased imbalance between AT2 and AT1 cells in β1^AT2-KO^ lungs throughout repair by calculating the number of pro-SP-C+ cells as a percentage of total cells (**Figure 3A**). Whereas the percentage of AT2 cells remained stable from day three into late repair in the β1^f/f^ lungs, the percentage of pro-SP-C+ AT2 cells progressively increased throughout later repair in β1^AT2-KO^ lungs. The increased AT2 number in β1^AT2-KO^ lungs prompted an examination of proliferation and apoptosis. We assessed AT2 proliferation by co-immunostaining lung sections for the proliferation marker Ki67 and pro-SP-C. β1^AT2-KO^ lungs exhibited increased proliferation in pro-SP-C+ AT2 cells at all sampled time points from uninjured lungs to day 21 post-injury (**Figure 3B**), with peak proliferation at day 7 (representative images shown in **Figure 3C**). We previously reported that β1 deficient AT2 cells exhibited increased NF-κB inflammatory signaling in the absence of injury (9). As NF-κB activation promotes cell survival and mitosis (23), we next tested whether the observed excess AT2 proliferation was due in part to upregulated NF-κB activation in β1 deficient AT2 cells. We measured AT2 proliferation in LPS-treated precision-cut lung slices (PCLS) generated from β1^AT2-KO^ and β1^f/f^ lungs in the presence/ absence of the NF-κB inhibitor BAY 11-7082. Proliferation was detected after 48 hours of treatment using incorporated BrdU, co-localized with pro-SP-C in immunostained PCLS. We confirmed induction of NF-κB activation with LPS *ex vivo* in PCLS by immunostaining for nuclear phospho-p65, signaling mediator in NF-κB activation (**Supp. Figure 5A**). Consistent with *in vivo* LPS dosing, β1^AT2-KO^ PCLS exhibited increased AT2 proliferation with LPS treatment (**Figure 3D-E**, individual values presented in **Supp. Figure 5B**), which was mitigated by concurrent NF-κB inhibition. For β1^f/f^ PCLS, AT2 proliferation rates remained similar in all four conditions. These data suggest that increased AT2 proliferation in β1^AT2-KO^ lungs post-injury is in part due to excess NF-κB activation intrinsic to β1-null AT2 cells.

**Figure 2.**
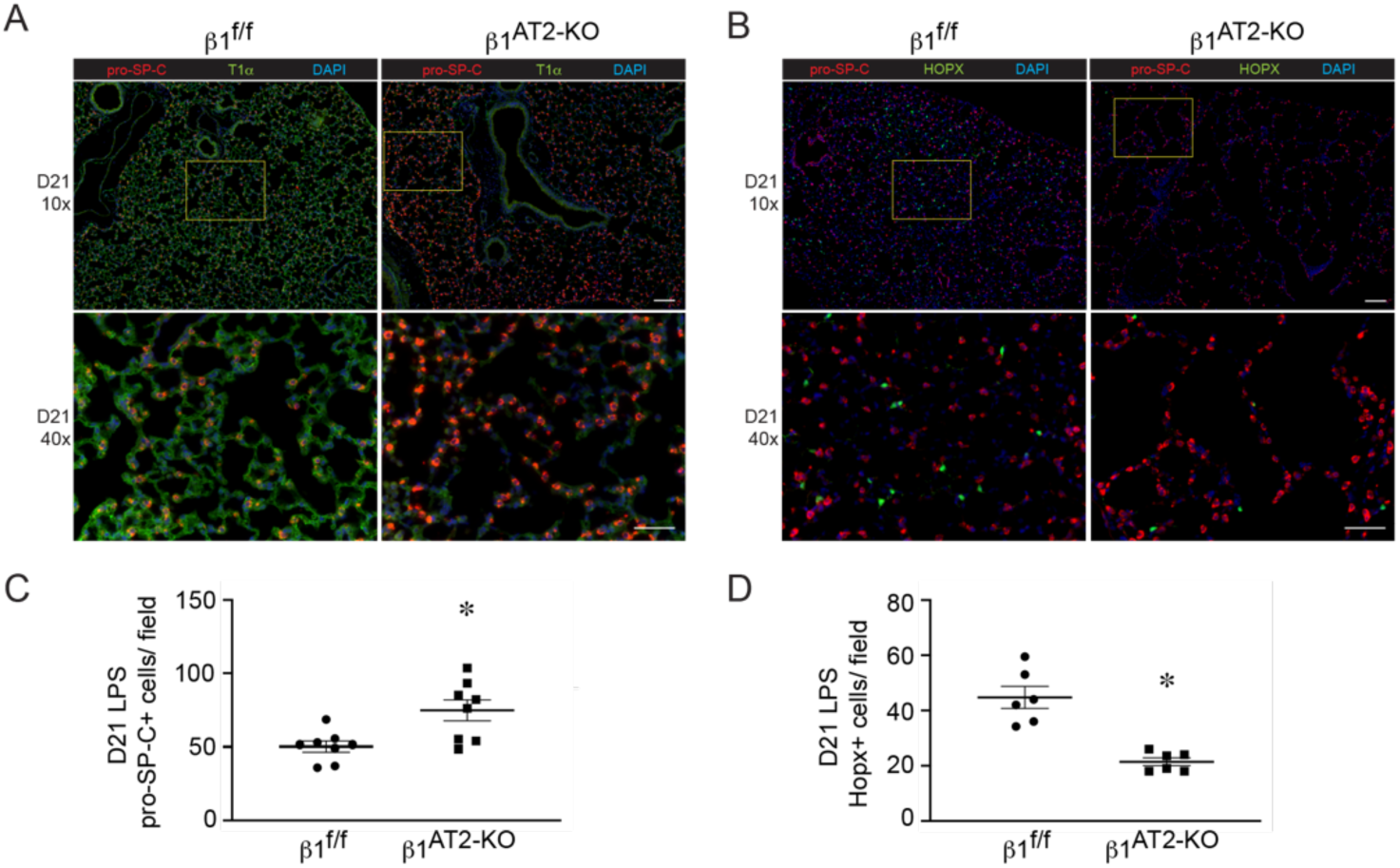
β1-deficient lungs have increased AT2 cell number and decreased AT1 cells at 21 days after LPS injury. (**A**) Representative low-power (10x) and inset high-power images (40x, denoted by yellow boxes) of β1^f/f^ and β1^AT2-KO^ lungs 21 days after LPS (D21), immunostained for AT2 marker pro-SP-C (red) and AT1 marker T1α (green). (**B**) Representative images of D21 β1^f/f^ and β1^AT2-KO^ lungs immunostained for pro-SP-C (red) with the AT1 marker HOPX (green), insets as in (A). (**C**) Quantification of the number of pro-SP-C+ AT2 cells per field (n=8 mice/ group, ten 40x sections/ mouse; *p*=0.0086). (**D**) HOPX+ cells per field (n=6 mice/ group, six 20x sections/ mouse; *p*=0.0003). * *p* < 0.05. Scale bar = 100 μm for 10x in A and B; scale bar = 50 μm for 40x in A and B. Two-tailed t-test was used to compare genotypes in C and D.

**Figure 3.**
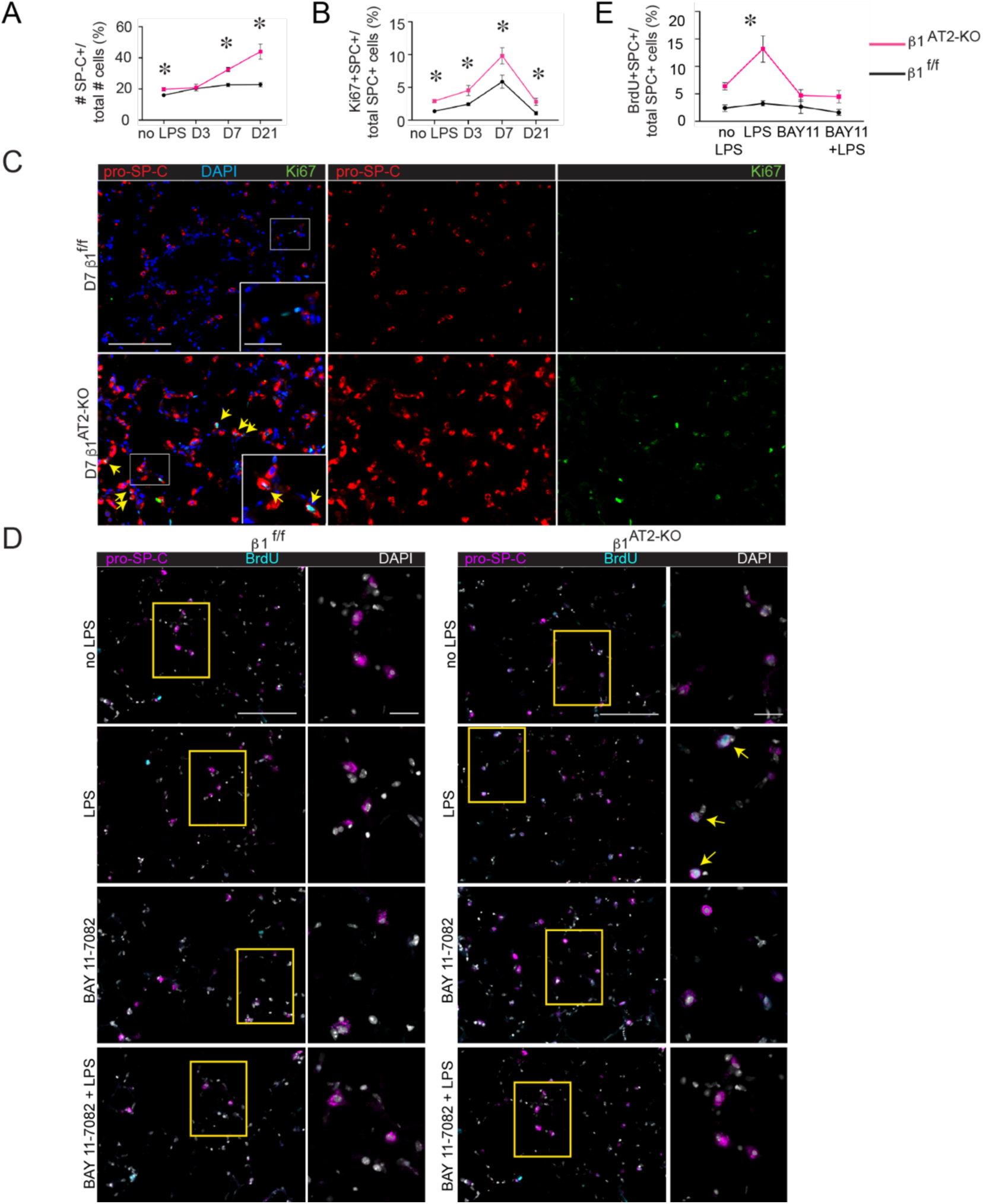
β1^AT2-KO^ mice have increased AT2 proliferation during alveolar repair. (**A**) Quantification of pro-SP-C+ cells per total cells (n=6-8 mice/ group; 5 sections per mouse; *p*=0.0247 for uninjured (no LPS) mice; *p*=0.8220 at D3; *p*=0.0001 at D7; *p*=0.0009 at D21). (**B**) Quantification of proliferating AT2 cells by percentage of total pro-SP-C+ AT2s (n=6-8 mice/ group, 10 sections/ mouse; *p*=0.0003 for uninjured mice; *p*=0.0311 at D3; *p*=0.0310 at D7; *p*=0.0128 at D21). (**C**) Immunodetection of the proliferation marker Ki67 (green) and pro-SP-C (red) show peak AT2 proliferation in β1^AT2-KO^ lungs at day 7 (arrows) as merged or single-channel panels. (**D**) Representative images of BrdU incorporated precision-cut lung slices (PCLS) treated with LPS and/ or NF-κB inhibitor BAY 11-7082 for 48 hours. Slices were immunostained for BrdU (cyan) and pro-SP-C (magenta) with DAPI nuclear marker (white). (**E**) Quantification of proliferating AT2 cells by percent of total AT2 cells (BrdU+pro-SP-C+ over total number of SP-C+ cells) by condition as indicated (n=6-8 mice/ group, 1 slice per mouse per condition, imaged and quantified ten 40x sections/ mouse per condition, data from five separate experiments; *p*=0.0010, F value=6.3 for treatment variation; *p*<0.0001, F value=26.1 for genotype variation). Two-tailed t-test was used to compare genotypes at each time point for A and B. Two-way ANOVA was used to compare treatment conditions and genotype in E. * *p* < 0.05. Scale bar = 200 μm for low-power in C and 50 μm for inset in C; scale bar = 100 μm for low-power in D and 50 μm for inset in D.

Analyzing cell survival by TUNEL assay co-immunostained with pro-SP-C, we identified a small but significant increase in the total number of apoptotic AT2 cells per field in β1^AT2-KO^ lungs on day 21 after LPS (**Supp. Figure 6A-C**). Although we were not able to capture large-scale AT2 apoptosis by TUNEL assay at these time points, early AT2 apoptosis remains a possibility. Further, the near-absence of AT1 cells seven days after injury strongly argues that AT1 apoptosis occurred early during acute LPS-induced injury.

### Overabundant AT2s are transcriptionally distinct during repair in β1^AT2-KO^ mice

To investigate the transcriptional phenotype of β1-deficient epithelial cells that could explain the development of emphysema and other features of abnormal repair in β1^AT2-KO^ mice, we performed single-cell RNA sequencing (scRNAseq) on unchallenged and LPS-treated β1^f/f^ and β1^AT2-KO^ lungs, seven days after LPS exposure. From single-cell suspensions from digested whole lung (n= 3-4 mice/ group pooled into a single sample for each condition), we collected CD45-/Ter119-viable cells by fluorescence-activated cell sorting (FACS), thereby excluding immune and red blood cells. We performed scRNAseq using the 10x Genomics platform, and doublet removal, normalization, and scaling led to 2,926 cells being analyzed. Initially, we integrated our dataset with recently published scRNAseq data from LPS injury (12), and we found similar alveolar epithelial cell types in both datasets (**Supp. Figure 7)**. To further define our epithelial populations throughout alveolar repair, we analyzed our transcripts utilizing the epithelial markers reported by Strunz et al with a label-transfer technique (11). In both LPS-treated and uninjured mice, we identified all major alveolar epithelial populations in both β1^AT2-KO^ and β1^f/f^ lungs (**Figure 4A**). Despite differential AT1 cell number difference in injured β1^AT2-KO^ lungs by histology, AT1 cells were isolated in low abundance in all four groups, uninjured and LPS-treated β1^AT2-KO^ and β1^f/f^ lungs, likely due to cell loss by FACS because of their relatively fragile structure (24). UMAP embedding identified a distinct epithelial population in β1^AT2-KO^ lungs seven days after injury (**Figure 4B**). Visualizing the UMAPs of the four individual groups, most of the epithelial cells were classified as AT2 cells or activated-AT2 cells, a cell state marked by inflammation-induced genes. In LPS-treated β1^AT2-KO^ lungs, the mixed AT2 and activated-AT2 cells were transcriptionally distinct from the other three groups. Because this defined β1^AT2-KO^ cluster comprised both AT2 and activated-AT2 cells, we combined these populations for further analysis in all four groups. We examined the top differentially expressed genes in the combined AT2 groups between uninjured and LPS-treated β1^AT2-KO^ or β1^f/f^ mice (**Figure 4C-D** and **Supp. Figure 8A-B**). In uninjured mice, the most enriched pathways were an upregulation of the oxidative stress-response pathway in the β1^AT2-KO^ AT2 group, followed by senescence, JAK/STAT, and IL-6 signaling pathway. All four of these top pathways being relevant to emphysema and lung senescence (25, 26), suggesting baseline conditions that favor accelerated aging in these three-month-old mice. In LPS-injured lungs from both genotypes, the top eight differentially up-regulated pathways included regulatory signaling networks related to adherens junctions, actin cytoskeleton, and Rho GTPases. As remodeling of adherens junctions is necessary for elongation of cell shape and Rho GTPases are intermediary signaling effectors between integrins and the actin cytoskeleton, these data suggest that β1 integrin regulates the transition in cell shape during AT1-cell differentiation.

**Figure 4.**
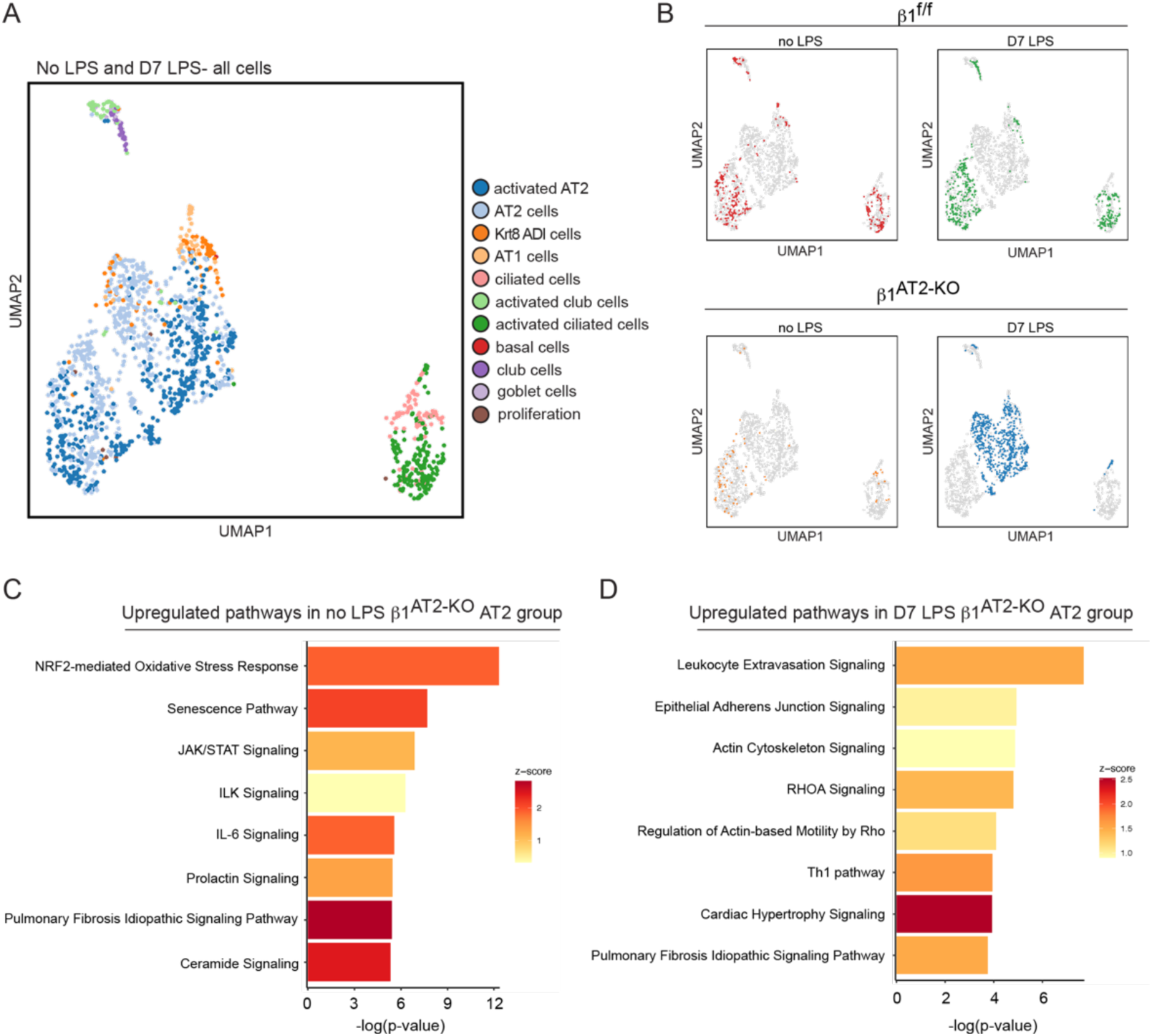
Overabundant AT2s are transcriptionally distinct during repair in β1^AT2-KO^ mice. (**A**) UMAP of all epithelial cells from β1^f/f^ and β1^AT2-KO^ lungs ± LPS clustered by label transfer from Strunz et al (11). (**B**) Individual epithelial populations by group reveal transcriptionally distinct AT2s and activated AT2s in day 7 LPS-treated β1^AT2-KO^ lungs. (**C**) Ingenuity Pathway Analysis on combined AT2 groups from uninjured β1^f/f^ and β1^AT2-KO^ lungs demonstrate upregulation of oxidative stress, senescence, and inflammatory pathways in β1^AT2-KO^ lungs compared to β1^f/f^ lungs. (**D**) Ingenuity Pathway Analysis shows upregulation of actin cytoskeleton signaling pathways in β1^AT2-KO^ AT2 cells compared to β1^f/f^ AT2 cells at seven days after LPS treatment.

### β1 integrin regulates actin expression and RhoA GTPase activation during alveolar repair

To investigate how β1 integrin regulates the actin cytoskeleton during alveolar repair, we examined AT2 cell shape by confocal microscopy. Surface rendering of pro-SP-C+ AT2 cells demonstrates a field of globular AT2 cells in β1^AT2-KO^ lungs, whereas β1^f/f^ pro-SP-C+ AT2 cells exhibited a flattened elongated cell shape (**Figure 5A**). We next imaged thick 50 μm cryosections detecting F-actin with phalloidin probe concurrent with pro-SP-C immunodetection. The injured β1^f/f^ pro-SP-C+ AT2 cells formed actin-based protrusions at the areas of their lateral extensions (arrows in **Figure 5B**), a feature shared with differentiating AT2 (to AT1) epithelial progenitors during development (27). By contrast, AT2s in injured β1^AT2-KO^ lungs were larger and rounded with cortical F-actin. We quantified AT2 cell shape at seven days post-injury by calculating cell area and a cell roundness score. Given the persistent macrophage inflammation during repair in LPS-treated β1^AT2-KO^ lungs, we specifically excluded the occasional macrophage that has efferocytosed an injured pro-SP-C+ cell from this analysis by calculating the day seven cell shape metrics on pro-SP-C+/ CD68-cells only (**Supp. Figure 9A**). AT2 cells were significantly larger in injured β1^AT2-KO^ lungs (66.8±3.0 μm^2^) compared to β1^f/f^ lungs (48.4±1.8 μm^2^) (**Figure 5C**). Since lateral extension is necessary for cells to transition from AT2 to AT1 and we demonstrated that cell shape is β1-dependent in AT2 cells, we calculated a roundness score, which quantifies the degree of smooth cellular contours in an unbiased manner (28). The roundness score in β1^AT2-KO^AT2 cells on day seven after injury, as they failed to elongate and change shape into flattened AT1 cells, was significantly increased compared to β1^f/f^ AT2 cells (**Figure 5D**). AT2 cells from LPS-treated β1^AT2-KO^ lungs maintained an elevated roundness score throughout repair, based on pro-SP-C+ cells (**Supp. Figure 9B**). This finding is consistent with an inability to form lateral extensions and impaired actin remodeling. By contrast, β1^f/f^ AT2 cell roundness decreased at times of rapid differentiation during repair, indicative of differentiation into a flattened AT1 cell shape. Since our scRNA-seq data suggested upregulation of actin-cytoskeleton signaling pathways, we next applied a G-actin probe, JLA20, along with F-actin phalloidin and pro-SP-C to study actin remodeling (**Figure 5E**). We found increased G-actin and F-actin in injured β1^AT2-KO^ lungs, with F-actin primarily cortically localized (arrowheads in Fig. 5E) in the β1^AT2-KO^ lungs. We confirmed these increases by calculating the relative fluorescence of the JLA20 and phalloidin probes in pro-SP-C+ cells using corrected total cell fluorescence to normalize fluorescence to total cell area (**Figure 5F-G**). Because actin remodeling and localization are mediated in part by the cytoskeletal protein ezrin, we immunodetected ezrin in β1^AT2-KO^ and β1^f/f^ lungs on day seven post-injury combined with *Sftpc* RNA *in situ* hybridization to mark AT2 cells. Ezrin was as expected localized to points of lateral cellular protrusion in elongating AT2 cells in control mice (**Figure 5H)**, while being much more widespread along the cell membrane in AT2s in β1^AT2-KO^ lungs, suggesting an impaired ability to direct actin localization in concert with loss of lateral elongation. We validated the linkage to small GTPases (29) suggested by scRNAseq analysis above using protein assays for specific GTPases. AT2 cells from LPS-injured β1^AT2-KO^ lungs exhibited increased RhoA and Cdc42 GTPase activation, as would be anticipated for a cell undergoing efficient actin remodeling (**Figure 5I-J**). By contrast, GTPase activation was decreased for Rac1 (**Figure 5K**), a related GTPase involved in forming smaller actin-based structures such as microvilli. Taken together, these data indicate RhoA and Cdc42 GTPase activation in AT2 cells is enhanced in β1 integrin deficient AT2 cells, as would be expected for a cell with active actin remodeling. However, evidence of actin remodeling at the site of lateral extension is absent, suggesting a block downstream of GTPase signaling events propagated by the β1 intracellular tail in β1^AT2-KO^ lungs. Thus, β1 integrin is a critical component for proper actin production and localization in differentiating AT2 cells during repair.

**Figure 5.**
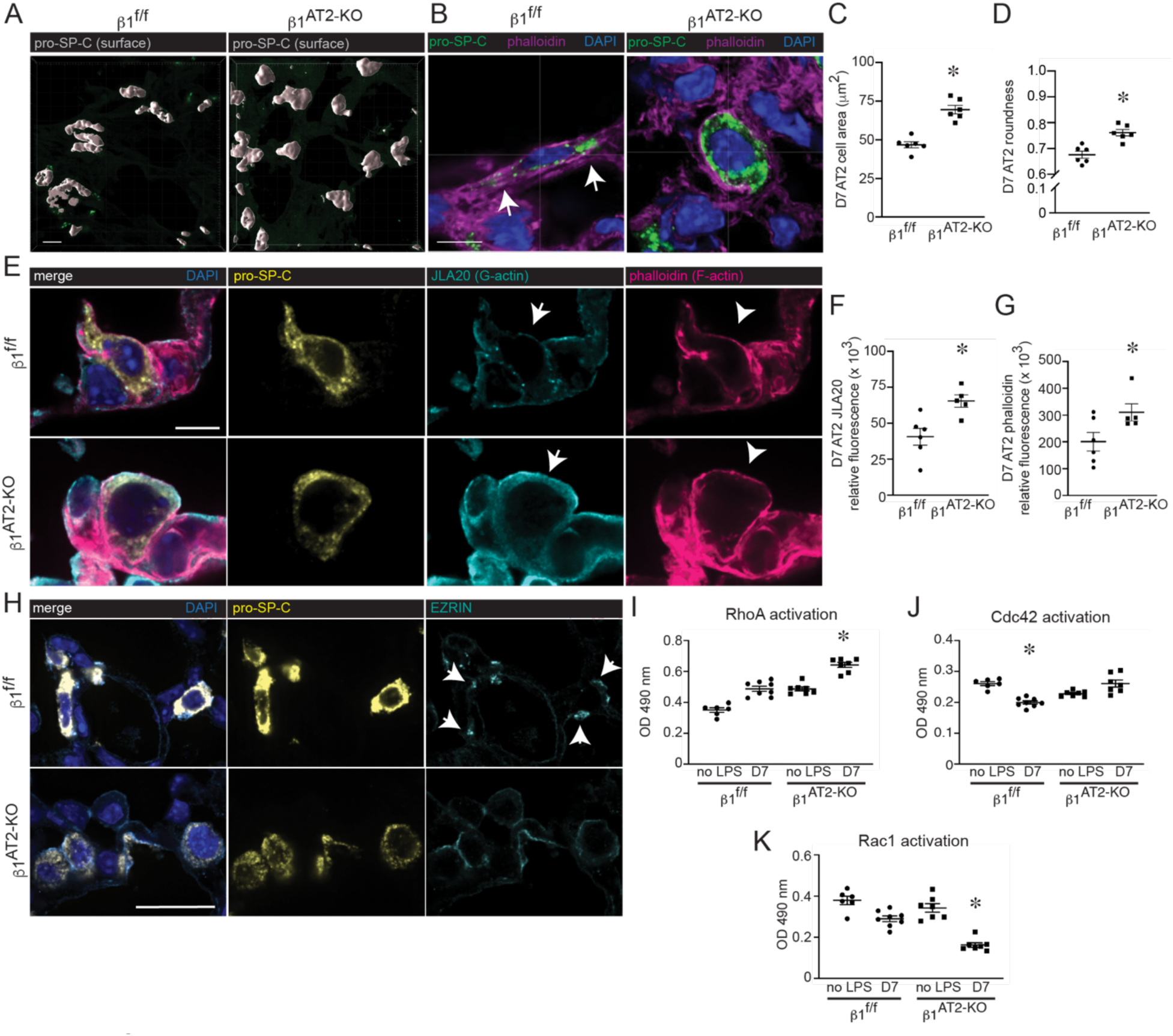
β1 integrin regulates actin localization and RhoA GTPase activation during alveolar repair. (**A**) Surface rendering high-power images of pro-SP-C immunostained thick frozen sections from day 7 (D7) LPS-treated β1^f/f^ and β1^AT2-KO^ lungs. (**B**) High-power images of thick frozen sections from D7 LPS-treated β1^f/f^ and β1^AT2-KO^ lungs immunostained for pro-SP-C (green) with phalloidin F-actin probe (magenta), arrows indicate areas of actin-rich lateral protrusions. (**C**) Area of pro-SP-C+/CD68-AT2 cells from D7 LPS-treated β1^f/f^ and β1^AT2-KO^ mice (46.8±2.0 μm^2^ in β1^f/f^ lungs compared to 69.6±2.8 μm^2^ in β1^AT2-KO^ lungs, n = 6 mice/group, ≥ 40 cells measured/ mouse imaged from 5 different sections, two-tailed t-test, *p*<0.0001). (**D**) Roundness score calculated from pro-SP-C+/CD68-cells from D7 LPS-treated β1^f/f^ and β1^AT2-KO^ mice (38-60 cells measured/ mouse from 5 different sections, n=6 mice/ group, two-tailed t-test comparing genotypes, *p*=0.0009). (**E**) High-power images of frozen sections prepared at D7 after LPS β1^f/f^ and β1^AT2-KO^ lungs immunostained for pro-SP-C (gold) with JLA20 (cyan) and phalloidin (magenta) probes applied to detect G-actin and F-actin, respectively. Membrane localization of G-actin denoted by arrows and F-actin by arrowheads. (**F-G**) Quantification of JLA20 (F) and phalloidin (G) expression in pro-SP-C+ AT2 cells in β1^f/f^ and β1^AT2-KO^ lungs D7 after LPS (n = 5-6 mice/ group, 10 sections/ mouse, two-tailed t-test with *p*=0.0088 for JLA20 and *p*=0.0482 for phalloidin). (**H**) Representative high-power images from day 7 LPS-treated β1^f/f^ and β1^AT2-KO^ lungs immunostained for ezrin (cyan) with AT2 cells identified by RNA *in situ* hybridization for *Sftpc* (gold). Arrows indicate ezrin expression localized to lateral extensions in β1^f/f^ AT2 cells, whereas diffuse, non-focal ezrin expression along the cell membrane is seen in β1^AT2-KO^ AT2 cells. (**I-K**) GTPase activation assay performed on AT2 cell lysates collected from uninjured and D7 LPS-treated β1^f/f^ and β1^AT2-KO^ lungs (n=6-8 mice/ group for each assay; RhoA one-way ANOVA *p*<0.0001, F value 53.42, df=3; Cdc42 one-way ANOVA *p*<0.001, F value 17.46, df=3; Rac1 one-way ANOVA *p*<0.0001, F value 31.09, df=3). * *p* < 0.05. Scale bar = 5 μm for A, B and E; scale bar = 25 μm for panels in H.

### Post-injury β1-deficient AT2s exhibit an AT2-AT1 mixed transcriptomic phenotype

Our observation of increased AT2 cells and decreased AT1 cells (Fig. 2) suggested impairment in AT2 to AT1 differentiation during alveolar repair. To define this defect at the transcriptional level we quantified lung epithelial cell sub-types as previously defined by Strunz et al (11). Consistent with our histology of the LPS-injured mice (Fig. 2), there was a significant increase in the representation of AT2 and activated AT2 cells in day seven post-LPS-injury β1^AT2-KO^ lungs, but not control β1^f/f^ (**Figure 6A**). We next compared the AT2, AT1, and cytokeratin 8–positive “intermediate cell” transcriptomic profiles in our uninjured and LPS-treated β1^AT2-KO^ and β1^f/f^ lungs. Strunz et al. (9) defined an intermediate cell state with features of both AT2 and AT1 cells as a “Krt8-positive alveolar differentiation intermediate (ADI)”. Both uninjured and LPS-treated β1^AT2-KO^ and β1^f/f^ AT2/activated-AT2 cells expressed AT2 hallmark genes (e.g., *Sftpc, S*ftpa1, and *Abca3*), as expected (**Figure 6B**). The Krt8+ ADI hallmark genes *Krt8, Hbegf,* and *Areg* were moderately increased in uninjured β1^AT2-KO^ AT2 cells and this increase was accentuated post-LPS. By contrast, there was minimal expression of these markers in uninjured or LPS-treated β1^f/f^ AT2 cells. Finally, uninjured β1^AT2-KO^ AT2/ activated AT2 cells exhibited enhanced AT1-cell marker expression compared to uninjured β1^f/f^ AT2 cells, which was also accentuated post-LPS. We validated increased AT1 marker expression in injured β1-deficient pro-SP-C+ AT2 cells by co-immunostaining for AGER, the AT1 marker with the highest expression by our transcriptional analysis. In injured β1^f/f^ lungs, AGER localized to the typical AT1 distribution throughout the lung parenchyma, largely distinct from pro-SP-C+ AT2 cells (**Figure 6C**, insets 1-3). By contrast, injured β1^AT2-KO^ lungs exhibited notably less AGER throughout the lung parenchyma compared to β1^f/f^ lungs. Importantly, the small amount of AGER present in β1^AT2-KO^ lungs co-localized with large, rounded pro-SP-C+ AT2 cells (insets 4-6, single channel images in **Supp. Figure 10**). The robust acquisition of AT1 markers in injured rounded, large β1^AT2-KO^ AT2 cells suggests that requisite β1 integrin-dependent mechanisms, such as cell shape change, occur subsequent to initial AT1 marker acquisition. In summary, absence of β1 increases the proportional representation of intermediate cell state markers, even in cells that remain classified as AT2 cells, indicating a significant block in the progression of differentiation.

**Figure 6.**
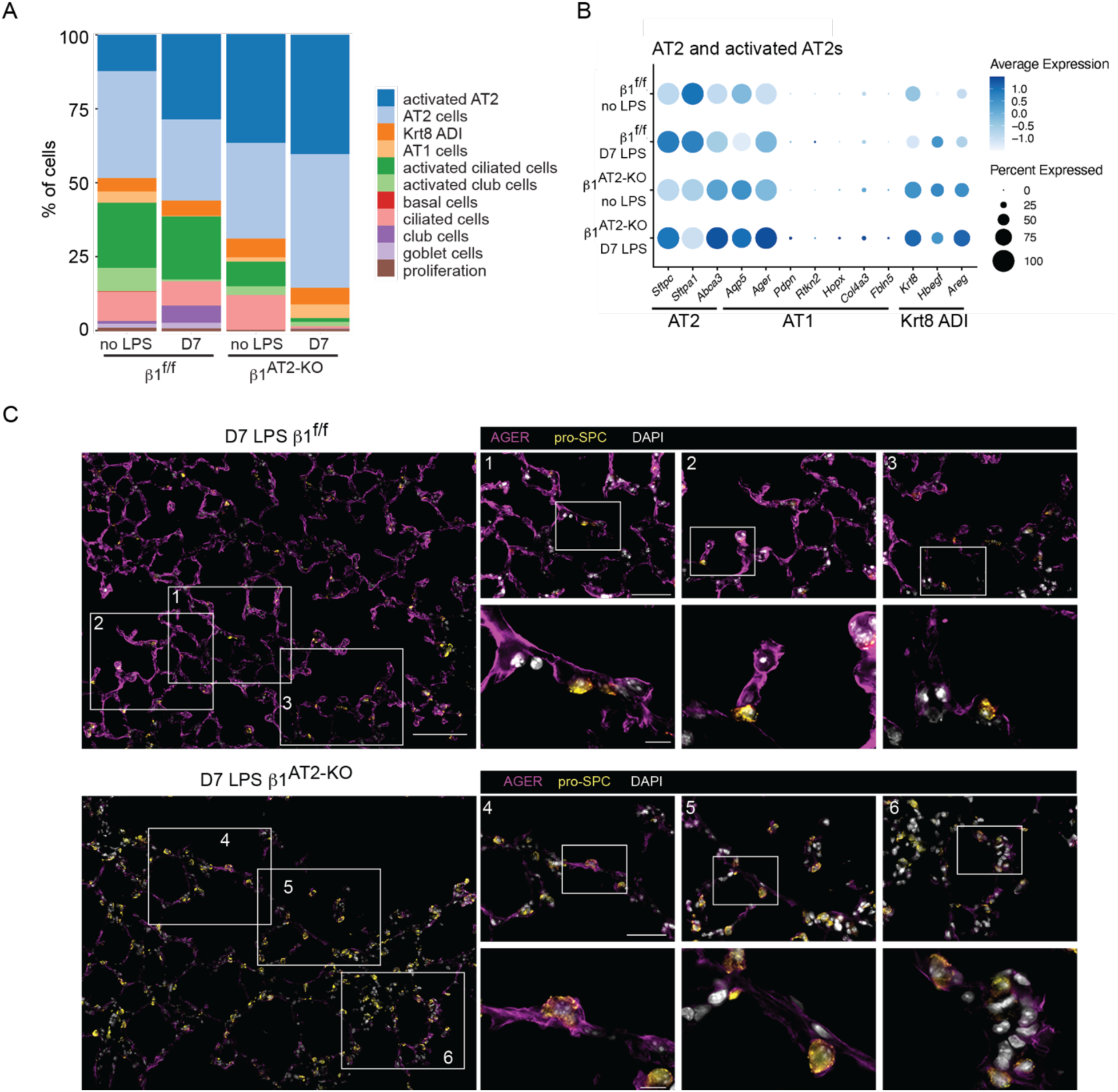
Post-injury β1 deficient AT2s exhibit an AT2-AT1 mixed epithelial transcriptomic phenotype. (**A**) Stacked bar graph of epithelial proportions demonstrates an expansion of the AT2 and activated AT2 populations in day 7 (D7) LPS-treated β1^AT2-KO^ lungs. (B) Marker gene expression by genotype and treatment group in AT2/ activated AT2 cluster, in which higher expression is represented with a darker color and the size of the dot reflects the proportion of cells expressing that marker. (**C**) Representative low-power images from day 7 LPS-treated lung sections immunostained for the AT1 marker AGER (purple) and AT2 marker pro-SP-C (gold) demonstrate overall decreased AGER in β1^AT2-KO^ lungs. Three insets per low-power field show co-localization of AGER with pro-SP-C+ AT2 cells in D7 β1^AT2-KO^ lungs. Scale bar = 100 μm for low-power in C, 50 μm for middle inset and 10 μm for high-power inset.

### Mixed-phenotype AT2 cells persist, proliferate, and maintain an enlarged, rounded cell shape in late alveolar repair

Since our histological and scRNA-seq analysis showed impaired AT2 differentiation at day 7 post-LPS, we characterized the blockage in differentiation in late repair. As described above (Fig. 3), we performed scRNA-sequencing on uninjured and LPS-treated β1^f/f^ and β1^AT2-KO^ lungs, 21 days post-LPS. Knowing that alveolar epithelial cells of mixed transcriptomic phenotype exist at day 7 post-LPS in β1^AT2-KO^ lungs, we investigated persistence of these cells at day 21 post-LPS by RNA sequencing of CD45-/Ter119-single-cell suspensions enriched for CD326+ epithelial cells. UMAP embedding with label transfer from Strunz et al. revealed similarities between β1^f/f^ and β1^AT2-KO^ alveolar cells (**Figure 7A-B**). By pro-SP-C/ Ki67 co-immunostaining, we demonstrated increased AT2 proliferation even during late repair (Figure 3B). Our scRNA-seq data confirmed that the AT2 population from day 21 LPS-treated β1^AT2-KO^ lungs exhibited a significantly increased G2M proliferation score compared to AT2 cells from day 21 LPS-treated β1^f/f^ lungs (**Supp. Figure 11A**). Similar to the scRNA-seq data from day seven, we next analyzed the expression of AT2, Krt8+ ADI, and AT1 hallmark genes in day 21 LPS-treated β1^AT2-KO^ and β1^f/f^ alveolar epithelial cells. We identified enhanced AT2 markers, *Sftpc*, *S*ftpa1, and *Abca3*, in AT2 and activated AT2 cell populations from β1^AT2-KO^ mice compared to β1^f/f^ mice (**Figure 7C**). In addition, β1^AT2-KO^ Krt8+ ADI cells retained AT2 hallmark gene expression compared to β1^f/f^ Krt8+ ADI cells, exhibited markedly enhanced expression of the three intermediate state markers (*Krt8, Hbegf,* and *Areg*), and increased expression of AT1 hallmark genes, *Aqp5* and *Ager*. These β1^AT2-KO^ Krt8+ ADI cells simultaneously expressed transcriptional markers from all three stages of transition, providing a transcriptional rationale for failed repair in β1-deficient mice. We validated the persistent mixed transcriptional phenotype in β1^AT2-KO^ lungs by immunodetection. In day 21 LPS-treated β1^AT2-KO^ lungs, remodeled areas exhibited numerous large, rounded pro-SP-C+/ cytokeratin 8+ cells with less AGER in the AT1 distribution compared to injured β1^f/f^ lungs (**Figure 7D**). High-power images demonstrate pro-SP-C+/ cytokeratin 8+ cells with occasional cells also positive for the AT1 marker AGER (arrows in **Figure 7E**). By contrast, triple positive cells (pro-SP-C+/ cytokeratin 8+/ AGER+) were rare in injured β1^f/f^ lungs (arrowheads in Figure 6E). The number of triple positive cells was quantified as a percent of total pro-SP-C+ cells in **Figure 7F**. Given the disparate cellular morphology of β1-deficient AT2 cells at earlier time points, we quantified differences in cell area in pro-SP-C+ AT2 cells. At 21 days after LPS, injured AT2 cells from β1^AT2-KO^ lungs remained enlarged with a significantly increased cell area (**Supp. Figure 11B**). These findings demonstrate that β1-mediated ECM interactions are essential during alveolar repair for maintenance of alveolar structure, epithelial cell shape change, and progression of their transcriptomic phenotype.

**Figure 7.**
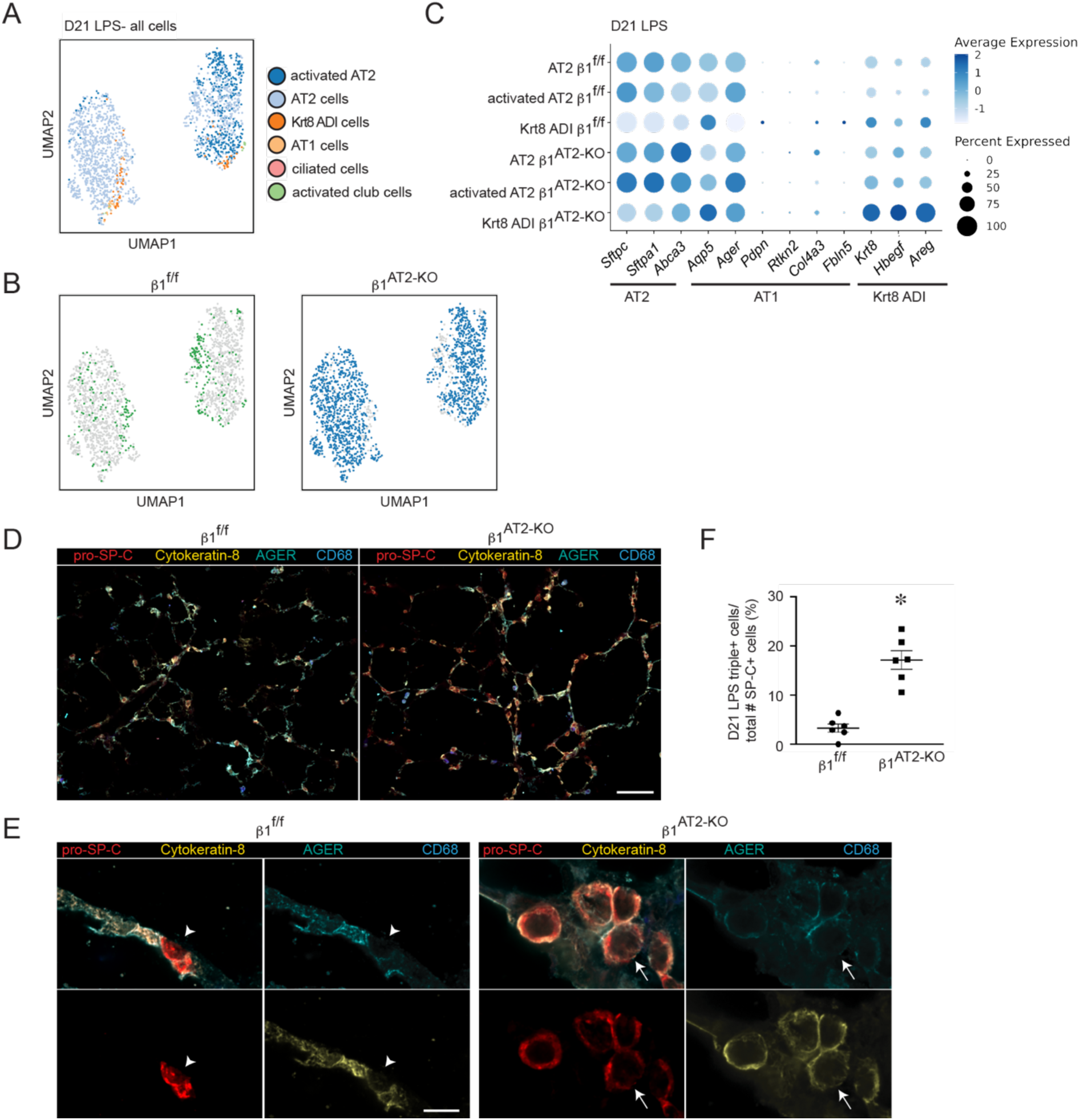
AT2s of mixed transcriptomic phenotype persist, proliferate, and maintain an enlarged, rounded cell shape in late alveolar repair. (**A**) UMAP of all epithelial cells from β1^f/f^ and β1^AT2-KO^ lungs 21 days after LPS treatment clustered by label transfer from Strunz et al (11). (**B**) Individual epithelial populations by group reveal transcriptionally abundant AT2s and activated AT2s in day 21 LPS-treated β1^AT2-KO^ lungs. (**C**) Hallmark gene expression by genotype in AT2, activated AT2, and Krt8 ADI clusters, in which higher expression is represented with a darker color and the size of the dot reflects the proportion of cells expressing that marker. (**D**) Representative low-power images of β1^f/f^ and β1^AT2-KO^ lungs 21 days after LPS co-immunostained for pro-SP-C (red), cytokeratin 8 (gold), AGER (cyan), and CD68 (blue). β1^AT2-KO^ lungs are notable for enlarged airspaces, increased numbers of round, large pro-SP-C+ AT2 cells, that are distinct from alveolar macrophages. (**E**) High-power images of lung sections immunostained for pro-SP-C, cytokeratin 8, AGER, and CD68, as above, demonstrate round, large pro-SP-C+ AT2 cells that co-localize with cytokeratin 8, with a subset also triple positive for pro-SP-C+/ cytokeratin 8+/ AGER+. Arrow denotes occasional triple+ cells in β1^AT2-KO^ lungs and arrowhead marks AGER-AT2 cell in β1^f/f^ lungs. (**F**) Quantification of pro-SP-C+/ cytokeratin 8+/ AGER+ triple positive cells as a percent of total pro-SP-C+ cells in day 21 LPS-treated lungs (n=6 mice/ group, ten 60x sections/ mouse, *p*<0.0001). * *p* < 0.05. Scale bar = 100 μm in D and 10 μm in E.

**Figure 8.**
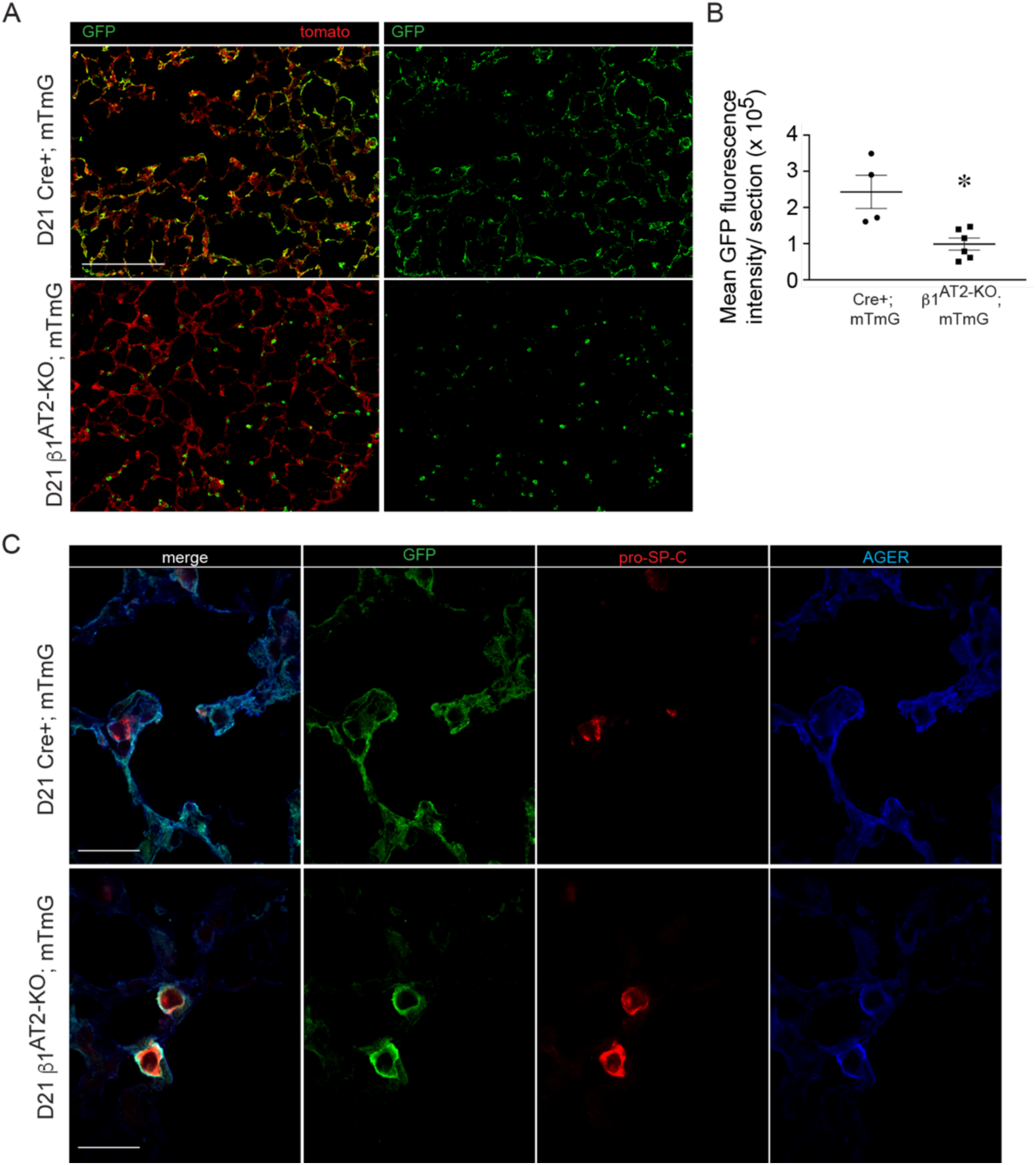
β1^AT2-KO^ mice fail to repopulate alveolar with cells of AT1 morphology. (**A**) Low-power images of Cre+; mTmG control mice demonstrate Cre-recombinase reporter labeled GFP positive cells of both AT1 and AT2 cell shape repopulate the injured alveolus 21 days after LPS. GFP labeled cells in β1^AT2-KO^; mTmG mice retain only AT2 morphology and fail to attain an AT1 cell shape at 21 days. (**B**) Quantification of mean GFP fluorescence intensity shows decreased area of lung repopulated by GFP labeled cells at 21 days post-LPS in β1^AT2-KO^; mTmG mice (n=4 Cre+; mTmG and 6 β1^AT2-KO^; mTmG mice; ten 20x sections/ mouse, *p*=0.0087). (**C**) High-power images of Cre+; mTmG and in β1^AT2-KO^; mTmG lung sections immunostained for pro-SP-C (red) and AGER (blue), tomato omitted in imaging. Large, round GFP+ labeled cells acquire both AT2 (pro-SP-C+) and AT1 (AGER+) markers in β1^AT2-KO^; mTmG lungs. GFP+ labeled cells possess either AT2 or AT1 markers at 21 days post-LPS in Cre+; mTmG lungs. * *p* < 0.05. Scale bar = 200 μm in A and 25 μm in C.

### β1-deficient AT2 cells fail to repopulate the alveolus with AT1 morphology

To prove that AT2 cells from β1^AT2-KO^ mice cannot repopulate the alveolus after injury, we crossed β1^AT2-KO^ mice with the mTmG Cre-recombinase reporter, labeling Cre-recombinase active AT2 cells beginning at P28. Mice were then maintained on doxycycline for a month and treated with LPS at three months-of-age. We confirmed the presence of GFP labeled AT2 cells prior to injury in β1^AT2-KO^; mTmG and Cre+; mTmG mice (**Supp. Figure 11A**). As anticipated, histological examination of lungs harvested 21 days after LPS demonstrates GFP positive cells in both AT2 and AT1 distributions in all the control Cre+; mTmG mice (**Figure 7A**), indicating patches of AT2 to AT1 alveolar repopulation post injury. By contrast, GFP labeled cells in β1^AT2-KO^; mTmG lungs retained almost exclusively a rounded AT2 morphology, with only rare acquisition of an elongated AT1 cell shape. We quantified the degree to which GFP labeled AT2 cells repopulated injured β1^AT2-KO^; mTmG and Cre+; mTmG lungs by fluorescence intensity per low field section (**Figure 7B** and **Supp. Figure 11B**). Immunostained sections for pro-SP-C and AGER confirmed the rounded GFP+ cells in β1^AT2-KO^; mTmG lungs were pro-SP-C+ and frequently AGER positive (**Figure 7C**). In Cre+; mTmG lungs, GFP labeling co-localized with AGER, confirming successful AT2 to AT1 differentiation in control mice. These data demonstrate failed AT2 to AT1 differentiation associated with impaired cell shape change and incomplete phenotypic epithelial marker differentiation in the absence of epithelial β1 integrin.

## Discussion

### β1 integrin is exquisitely required for differentiation during repair, in contrast to development or alveolar homeostasis

Since progenitor AT2 proliferation precedes differentiation in the injury-repair cycle, the overabundance of incompletely differentiated AT2 in β1^AT2-KO^ mice reveals a substantial block in differentiation post-injury. We previously reported increased AT2 proliferation with deletion of epithelial β1 in both the developing and adult lung in the absence of injury (8, 9). The degree of AT2 accumulation in β1-deficient repair reported here far exceeds the developmental and homeostatic contexts. This relative difference in AT2 surplus, coupled with successful AT2 to AT1 differentiation in uninjured aged β1^AT2-KO^ mice (9), shows that β1-ECM interactions are even more critical for differentiation during repair of adult injury than during development or homeostasis. This comparison suggests that β1-dependent differentiation mechanisms are specifically adjusted to fit the requirements of the peripheral stem-cell niche. One possible explanation is that the time interval over which cells change shape and differentiate differs between the three contexts of development, homeostasis, and repair. Developmental AT2 to AT1 differentiation occurs over weeks, and sporadic AT2 replacement of AT1 cells in the adult lung occurs without the urgency of replacing a widely denuded BM. By contrast, AT2 differentiation post-LPS happens en masse with the BM exposed within hours and repopulated within days. This shorter time interval makes robust upregulation of facultative differentiation pathways less likely, exposing a dependence on β1-mediated mechanisms for regulating differentiation. Another possible reason for the β1 requirement during repair may be related to the condition of the BM. It is well known that changes in the relative amounts of ECM components will push the alveolar epithelium towards proliferation or differentiation (30), although less is known about how structural changes to the BM itself drives epithelial behavior in the alveolar stem cell niche. The mature uninjured lung has a relatively stable BM with slow turnover of BM proteins (31). However, during LPS-induced injury, inflammatory-cell–associated proteases partially degrade the alveolar BM within hours (3, 4, 32). Numerous proteases degrade ECM components preferentially, thereby significantly modifying the carefully balanced alveolar BM composition (4). As different injuries induce varying types of protease-release, damage to the BM components will change depending on type and severity of insult, further shifting β1-ECM interactions in the stem-cell niche away from homeostatic conditions. In addition, individual β1-containing integrin pairs preferentially bind their preferred ECM ligands, thereby setting up a situation where injury-induced changes to β1-ECM interactions can either restrict or augment epithelial differentiation.

The progressive increase in AT2 cell number that escalates late in repair in β1^AT2-KO^ lungs indicates that β1-ECM interactions may serve as a brake on AT2 progenitor cell proliferation, could be a requisite step for progression from proliferation to differentiation during the alveolar injury repair cycle, and that this stalled differentiation stimulates a feed-forward loop that creates a continued abnormal call for progenitor proliferation. The lung is unique among branched organs in that there exists a distal (in fact, intra-alveolar) epithelial progenitor representing an indigenous contributor to tissue regeneration (13–15, 33), whereas no such epithelial progenitor has been reported for the kidney, mammary gland, or pancreas. Activation of the NF-κB pathway is pro-survival and pro-proliferative in many contexts, initially reported in hyperproliferative keratinocytes with constitutive NF-κB activation (23). While multiple signaling pathways have been proposed as positive and negative regulators of AT2 progenitor proliferation in the adult lung (13-16, 34-37), their interaction with the NF-κB pathway in the injured lung remains largely undefined. Although a recent study by Sieber et. al reports NF-κB-driven airway proliferation in human IPF airway epithelial/ fibroblast co-cultures (38). In other organs, β1-containing integrins have been reported to negatively regulate epithelial proliferation, with an EGFR-activation dependent mechanism identified in the intestinal epithelium (39), but these studies do not consider progenitor proliferation in the context of repair. Consideration of our data in light of this published work suggests that β1 integrin restricts proliferative potential in progenitor AT2s in the injured distal lung.

### Failed differentiation in injured β1-deficient mice reveals that cell shape change is intimately tied to progression of differentiation

We demonstrate that β1-ECM interactions are a requisite component for epithelial cell shape change through actin remodeling in the injured alveolus. The connection between integrins and cellular extension comes from studies in Drosophila epithelium and murine podocytes, both in the *uninjured* state (40–44). Our scRNA-sequencing analysis supports the concept that cell shape change is linked to progression of differentiation, as actin remodeling pathways were enhanced in LPS-treated β1-deficient AT2 cells, but differentiation remained blocked with an overabundance of large, rounded AT2 cells. It is possible that loss of integrin clustering at the cell membrane disrupts the spatial specificity of Rho-GTPase signaling (45). Certainly, diffuse ezrin expression associated with persistence of a rounded cell shape and loss of lateral cellular extensions observed in LPS-treated β1^AT2-KO^ AT2 cells is consistent with impaired actin nucleation.

Historically, cellular identity has been defined by histological shape and directly observable characteristics of function (e.g., lamellar body synthesis by electron microscopy). The advent of single-cell transcriptomics increases the granularity with which we can group similar cells together. While this advance augments our understanding of cellular transitional states, the question of how to define groups of similar cells and the boundary between different cell identities is perhaps murkier. Previously published injury models report similar temporary and resolvable transitional cell states after bleomycin injury and other inflammatory stimuli (17, 18), which are marked by high *Krt8* expression. Our transcriptomic data demonstrated persistently blocked AT2-to-AT1 differentiation in injured β1^AT2-KO^ lungs, suggesting that they are stuck in a transitional state even in late repair. Although our model is unique in the degree and persistence of failed differentiation, our transcriptomic data has some similarities and differences with recently published single-cell sequencing data from LPS injury (12). The observed differences in the transitional cell-state populations identified are possibly linked to technical differences in how epithelial cells were sorted; Riemondy et al. used AT2 cells selected by expression of a fluorescent reporter for *Sftpc*-expression prior to scRNAseq, removing transitional cells with lower *Sftpc* expression.

### Loss of β1-mediated epithelial-ECM interactions increases susceptibility for emphysematous abnormal repair and accelerates aging

Loss of β1 integrin in AT2 cells generates a senescent transcriptional signature in an otherwise uninjured youthful lung that translates into emphysematous abnormal repair with injury. Emphysema is a principal component of chronic obstructive pulmonary disease (COPD), one of the most common human pulmonary diseases that increases in incidence with age (46, 47). Although airspace expansion has been reported with either two-hit injury models or chronic repetitive low-dose LPS exposure (48–52), our β1-deficient mice are the first targeted epithelial deletion that results in subacute alveolar destruction following a single LPS exposure, underscoring the importance of epithelial β1 integrin in senescence and its associated pulmonary pathologies. Our scRNA-sequencing pathway analysis demonstrates uninjured β1-deficient AT2 cells possess an inherent susceptibility to injury, as uninjured β1^AT2-KO^ mice exhibit upregulation of oxidative stress, senescence, and inflammatory pathways, all contributing factors to emphysema pathogenesis. LPS-challenged β1^AT2-KO^ mice also exhibit increased inflammation during repair, an associated finding in COPD (53), with a sustained increase in neutrophils at day 7 after LPS, potentially making prolonged exposure to elastase a contributing factor for emphysema development. At 21 days post-injury, β1^AT2-KO^ lungs retain elevated numbers of macrophages, a significant source of matrix metalloproteinase-9 (MMP-9), which remodels alveolar matrix in COPD (50). It is unlikely that increased AT2 apoptosis in β1^AT2-KO^ lungs is the primary mechanism for the development of emphysema as it was only present late in repair at day 21, when emphysematous structural changes had already occurred. The etiology for emphysematous alveolar remodeling in young β1 deficient mice is likely multi-factorial, but rooted in the preexisting upregulation of senescence, oxidative stress, and inflammatory pathways. Additional studies addressing how β1 integrin regulates these pathways would elucidate if and how β1-dependent mechanisms could increase susceptibility of human COPD.

In conclusion, this study shows that β1 integrin-mediated mechanisms are a requisite component of rapid and complete alveolar repair following even mild LPS-induced lung injury, and involve regulation of AT2 proliferation, differentiation, and cell-shape change. In the latter, we propose that cytoarchitectural changes, such as the formation of actin-rich lateral cellular extensions, are specifically tied to progression of AT2 to AT1 differentiation during repair. Loss of β1 integrin results in upregulation of senescence, oxidative stress and inflammatory pathways already pre-injury, thereby increasing susceptibility to emphysema, a common age-related lung structural deficit.

## Methods

### Mice and LPS injury

We induced β1 integrin in AT2 cells in the adult murine lung by crossing transgenic mice with inducible Cre recombinase expression by the doxycycline-inducible reverse tetracycline transactivator under control of the SP-C promoter (SP-C rtTA; Tet-O-Cre) with integrin β1^flox/flox^ (β1^f/f^) mice (54, 55). Littermate rtTA negative; Tet-O-Cre negative (β1^f/f^ mice were used as control mice. Doxycycline drinking water (2 g/L) was administered for 4 weeks beginning at P28 to triple transgenic SP-C rtTA; Tet-O-Cre; β1^f/f^ mice (called β1^AT2-KO^ mice) and littermate β1^f/f^ controls, as previously described (9). Following proposed guidelines to limit Cre leak with the doxycycline-inducible system (20), we used the more faithful line 2 strain, employed a breeding strategy to keep the rtTA allele hemizygous, and utilized an injury model that targets the peripheral lung rather than the airway epithelium. After a 1-month period off doxycycline to minimize any potential toxicity from tissue stored doxycycline, we challenged 3-month-old β1^AT2-KO^ and β1^f/f^ mice with a single intratracheal dose of LPS or PBS (3 μg/g mouse weight, equivalent volume for PBS) and harvested tissue at the indicated dates. For β1^AT2-KO^ and β1^f/f^ mice administered delayed doxycycline, intratracheal LPS was given at three months of age as above, then doxycycline drinking water was provided from five days after LPS until harvest at 21 days after LPS. SP-C rtTA; Tet-O-Cre (no floxed alleles) and β1^AT2-KO^ mice were crossed to the mTmG Cre recombinase reporter. Doxycycline drinking water and LPS were administered as above. SP-C rtTA, Tet-O-Cre, and mTmG mice were purchased from Jackson Laboratory. Integrin β1^f/f^ mice were generously gifted by Elaine Fuchs (Howard Hughes Medical Institute, The Rockefeller University, New York, NY). All mice were maintained on the C57BL/6 background.

### Histology and morphological analysis

For histological analysis on paraffin sections, mice were sacrificed, right ventricle was flushed with PBS, and lungs were inflation fixed at 25 cm with 10% formalin. After paraffin processing, embedding, and sectioning, lungs were hematoxylin and eosin stained for morphological analysis by mean linear intercept, which was calculated from images (≥ 10 non-overlapping images per mouse) obtained using a x40 objective on a Keyence BZ-X710 inverted fluorescence phase contrast microscope. Immunofluorescence staining was performed on paraffin or frozen sections. Frozen blocks were prepared from lung sections inflation fixed with a 2:1 PBS/O.C.T. (Tissue-Tek) mixture, embedded, and sectioned at either 8 μm or 50 μm thickness. Frozen section slides were then fixed with 4% paraformaldehyde, permeabilized with .1% Triton-X, and blocked with 5% donkey serum for 2 hours at 37 degrees. Slides were incubated in primary antibody overnight at 4 degrees, followed by secondary antibody incubation for 2 hours at room temperature. For sequential primary antibody staining, a second intervening blocking step was utilized. Nuclei were stained with Dapi, ProLong Gold mountant was applied, and sections were imaged. High-power images were obtained using a Nikon Spinning Disk TiE inverted fluorescent confocal microscope attached to an Andor DU-897 EMCCD camera (x100 objective, 8 μm sections). All other images were obtained using the Keyence BZ-X710 microscope as above. The following primary antibodies and probes were used: anti-pro-SP-C (Abcam ab90716), anti-ABCA3 (Abcam ab24751), anti-T1α (podoplanin, Developmental Studies Hybridoma Bank 8.1.1), anti-ezrin (Cell Signaling Technologies 3145S), anti-cytokeratin 8 (Origene BP5074), anti-Ki67-FITC (eBioscience 11-5698-90), anti-CD68 (Abcam ab53444 and ab125212), anti-Ager (R and D AF1145), anti-Hopx-647 (Santa Cruz sc-398703), anti-BrdU (Sigma B2531), anti-GFP (Abcam ab13970), anti-pan-laminin (Millipore AB2034), anti-integrin β1 (Millipore MAB1997), anti-integrin α5 (Abcam ab150361), anti-integrin αv (Abcam ab179475), anti-integrin β5 (Invitrogen PA5-118499), anti-integrin β6 (Invitrogen PA5-47309), anti-integrin β8 (Invitrogen PA5-100843), anti-phospho-p65 (Abcam ab194726), JLA20 (Developmental Studies Hybridoma Bank JLA20-s), and phalloidin (Invitrogen A12380). We applied the following secondary antibodies: anti-rabbit Alexa Fluor 405 (Invitrogen A48258), anti-rabbit Alexa Fluor 488 (Invitrogen A21206), anti-rabbit Alexa Fluor 594 (Invitrogen A21207), anti-rabbit Alexa Fluor 647 (Invitrogen A32795), anti-rat Alexa Fluor 405 (Invitrogen A48268), anti-hamster Alexa Fluor 488 (Invitrogen A21110), anti-goat Alexa Fluor 488 (Invitrogen A11055), anti-goat Alexa Fluor 647 (Invitrogen A21447), anti-chicken Alexa Fluor 488 (Invitrogen A11039), anti-guinea pig Alexa Fluor 647 (Invitrogen A21450). TUNEL staining was performed on paraffin sections co-immunostained with pro-SP-C, per manufacturer’s kit instructions (Roche 11684795910). RNA *in situ* hybridization for mouse *Sftpc* was used in conjunction with immunofluorescent staining for the anti-ezrin antibody. RNA *in situ* hybridization was performed per manufacturer’s instructions (RNAScope, ACDBio), including positive and negative control probes. Quantification of immunostained sections was performed on ≥ 10 nonoverlapping images obtained with a 40x objective. JLA20 and phalloidin probe detection was quantified on lung sections (10 sections/ mouse) imaged with equivalent settings using the corrected total cell fluorescence feature from ImageJ, which corrects fluorescence integrated density for the area of the region of interest (pro-SPC+ cells).

### Cell morphometry

Day 7 and day 21 AT2 cell area and roundness were calculated using the shape descriptor feature of ImageJ with pro-SP-C+/CD68-cells used to define the region of interest.

### Precision-cut lung slices (PCLS)

PCLS of 300 μm were generated from the lungs of 3-month-old β1^AT2-KO^ and β1^f/f^ mice, as previously described (56–58). Briefly, Slices were cultured in Dulbecco’s modified Eagle medium: F12 for 12 hours, then incubated with BrdU (1 mM, Sigma B5002) for 4 hours prior to 48 hours of treatment with LPS (Sigma L2880, 62.5 ng/ml) and/ or BAY 11-7082 (100 μM, Biotechne/Tocris Bay11-7821). Slices were then fixed, paraffin-embedded, and sectioned.

### Bronchoalveolar lavage

After sacrifice, lungs were lavaged with 1 ml sterile PBS. The cells present in lavage fluid were collected by centrifugation, resuspended, and counted. Protein content in lavage fluid was measured by BCA protein assay (Pierce cat #23225) per manufacturer’s instructions.

### AEC isolation and GLISA assay

Primary AT2 cells were collected at indicated time points as previously described (8, 9), which yields > 90% AT2s (59, 60). Briefly, single cell lung suspension was generated after dispase digestion and serial filtration. Cells were then applied to plates coated with CD-32 and CD-45 for negative selection. Epithelial cells were then collected from medium after a 2 hour incubation. Cell lysates were then used for G-LISA small GTPase activation assay (Cytoskeleton cat# BK135), where levels of activated RhoA, Cdc42, and Rac1 were detected colorimetrically per manufacturer’s instructions.

### Single-cell data collection

Sample collection and single-cell sequencing was performed as previously described (61). Briefly, lung lobes were harvested, minced, and incubated for 30 minutes at 37°C in dissociation media (RPMI-1640 with 0.7 mg/ml collagenase XI and 30 mg/ml type IV bovine pancreatic DNase). After incubation, tissue was disassociated into a single-cell suspension by passage through a wide bore pipet tip and filtration through a 40 µm filter. The single-cell lung suspension was then counted, aliquoted, and blocked with CD-32 Fc block (BD cat # 553142) for 20 minutes on ice. After a 2% FBS staining buffer wash, cells were incubated with the conjugated primary antibodies anti-CD45 (BD cat # 559864) and anti-Ter119 (Biolegend cat# 116211). Samples from day 21 after LPS were also incubated with anti-CD326 antibody (BD cat # 563477) for epithelial enrichment.

### scRNA-seq library preparation and next-generation sequencing

scRNA-seq libraries were generated using the Chromium Single Cell 5’ library preparation kits (10X Genomics) following the manufacturer’s recommendations and targeting 10,000 - 20,000 cells per sample. Sequencing was performed on an Illumina Novaseq 6000. CellRanger Count v3.1 (10X Genomics) was used to align reads onto the mm10 reference genome.

### Analysis of single cell sequencing data

Ambient background RNA was cleaned from the scRNA-seq data with “SoupX”(62) as described previously (61) using the following genes to estimate the non-expressing cells, calculate the contamination fraction, and adjust the gene expression counts: *Dcn*, *Bgn*, *Aspn*, *Ecm2*, *Fos*, *Hbb-bs*, *Hbb-bt*, *Hba-a1*, *Hba-a2*, *Lyz1*, *Lyz2*, *Mgp*, *Postn*, *Scgb1a1*. For all datasets, quality filtering was then used to remove cells with > 15% or < 0.1% mitochondrial mRNA, and to remove cells with < 700 detected genes.

Dimensionality reduction, clustering, and visualization was performed using Seurat v4.0.5 and SCTransform v0.3.2.9008 with glmGamPoi v 1.6.0 (63–65). SCTransform was run with each sequencing run as a batch variable, and with the percentage of mitochondrial RNA as a regression variable. Further data cleaning was done to remove gene counts for *Gm42418*, which is likely a rRNA (66). Epithelial cells (*i.e., Epcam+* cell clusters) were sorted *in silico* for downstream analysis. Epithelial cells were annotated with manual inspection of the following marker genes: *Epcam*, Sftpa1, *Sftpc*, *Hopx*, *Aqp5*, *Col4a3*, *Ager*, *Foxj1*, *Dynlrb2*, *Mki67*, *Scgb1a1*, *Scgb3a2*, *Cdkn1a*, *Cldn4*, *Ascl1*, and *Scg5*. All charts and heatmaps as part of the scRNAseq analysis were generated with ggplot2, and all parts of the analysis were run in R 4.1.1.

Data integration was performed using a previously published dataset (12). Reads were downloaded from the Gene Expression Omnibus (GSE113049) and processed using CellRanger Count v3.1 with the mm10 genome, followed by dimensionality reduction, clustering, and visualization as described above. Data were integrated with non-SoupX processed data using the ‘IntegrateData’ function in Seurat, following the workflow for data normalized with SCTransform. Samples from all experiments were combined, and clusters were annotated using marker genes described in the initial publication (12), and other canonical marker genes.

Cell label transfer was also utilized with a second published dataset (11). The annotated data matrix (h5ad file) was accessed as directed in the analysis code published as part of the methods (11). The raw counts from this annotated dataset were then processed using the Seurat workflow described above, while maintaining the published cell annotations. The Seurat “TransferData” function was then used to transfer the cell annotations onto the dataset generated here.

A complete collection of all package versions, and code for all steps of the analysis is available at https://github.com/SucreLab. All sequencing data have been deposited to the NCBI GEO database; the reviewer link is available upon request.

### Statistics

Comparison between two groups was performed by two-tailed t-test and 4-way comparison was done by one or two-way ANOVA, as indicated. Individual *p-*values, t-values, F values, degrees of freedom, and sample size for each group is included in the figure legends and provided in **Supp. Table 1**. ScRNA-sequencing analysis was completed as above.

### Study approval

All animal experiments were approved by the Institutional Animal Care and Use Committee at Vanderbilt University Medical Center.

## Author contributions

EJP conceived the study, performed in vivo experiments, histological analysis, image analysis, interpreted the data, and wrote the manuscript. JMSS interpreted the scRNA-sequencing data and wrote the manuscript. FB, JG, JTB, and KTF performed histological analysis. PMG performed in vivo experiments, histological analysis, and protein assays. XD and SK performed protein assays. WH, CSJ performed in vivo experiments. JAK, NMN, and YL analyzed the scRNA-sequencing data. SHG assisted in manuscript preparation. RZ and TSB conceived of the study, interpreted the data, and wrote the manuscript.

## Acknowledgements

The authors would like to thank Matt Tyska and Christopher V.E. Wright for content expertise and Brittany Matlock, Angela Jones, Kari Seedle, Caitlin McCormick, Cristina Harmelink, Annica Alwine, and David Nichols for technical assistance. This work was supported by NIH grants K08 HL127102 (EJP), R03 HL 154287 (EJP), R01 HL163195 (EJP), K08 HL143051 (JMSS), P01 HL092870 (TSB), R01 HL151016 (TSB), R01 DK069921 (RZ), R01 DK127589 (RZ), R01 DK088327 (RZ), R01 HL157373 (JTB), R01 HL150617 (SHG), R01 HL153246 (JAK), R38 HL143619 (trainee: KTF), Merit Review Grant 2 I01 BX002378 (TSB), I01 BX002196 (RZ), the Parker B. Francis Family Foundation (JMSS, JJG), American Society of Nephrology Ben J. Lipps Research Fellowship (FB), and the Vanderbilt Faculty Research Scholars award (JJG). Experiments were performed utilizing the Vanderbilt Cell Imaging Shared Resource (supported by NIH grants CA68485, DK20593, DK58404, DK59637, and EY08126). ScRNA-sequencing experiments were performed using the Vanderbilt Technologies for Advanced Genomics Core facility. Flow cytometry experiments were performed in the VUMC Flow Cytometry Shared Resource (supported by the Vanderbilt Ingram Cancer Center P30 CA68485 and the Vanderbilt Digestive Disease Research Center DK058404).

