## Supplemental Materials for "Alveolar Repair Following Lipopolysaccharide-induced Injury Requires Cell-Extracellular Matrix Interactions"

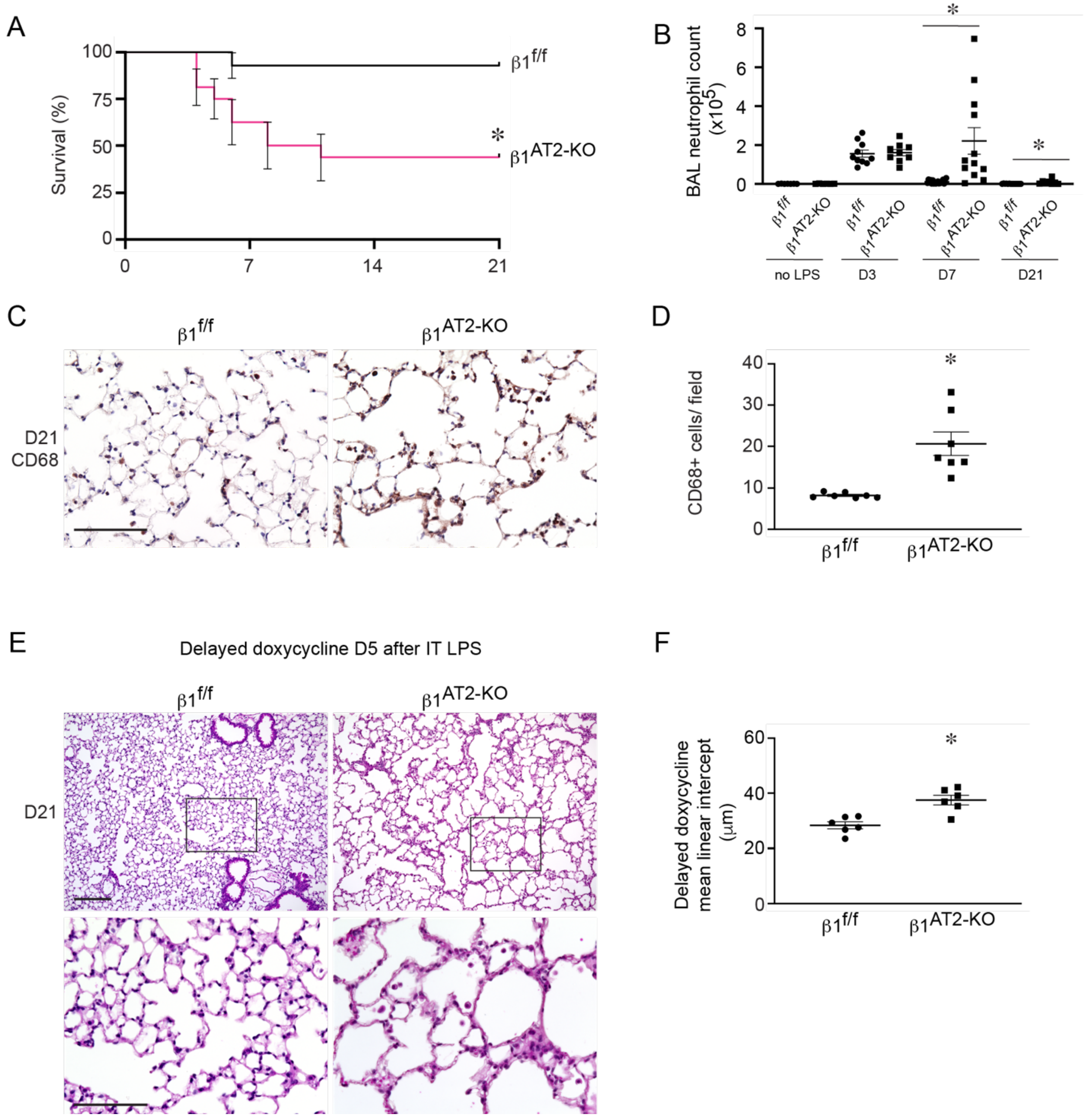


**Supplemental Figure 1. Emphysema in $\beta1^{AT2\text{-}KO}$ mice develops during alveolar repair, independent of acute inflammation.** (**A**) Kaplan-Meier 21-day survival curve following intratracheal LPS administration demonstrates significantly decreased survival in $\beta1^{AT2\text{-}KO}$ mice compared to control $\beta1^{f/f}$ littermates (44% survival in $\beta1^{AT2\text{-}KO}$ mice compared to 93% in $\beta1^{f/f}$ mice, n=15 $\beta1^{AT2\text{-}KO}$ and 14 $\beta1^{f/f}$ mice, * $p$=0.0051, Chi square 7.855 by log-rank test). (**B**) Neutrophil counts are increased at day 7 and day 21 in $\beta1^{AT2\text{-}KO}$ BAL compared to $\beta1^{f/f}$ BAL (n = 6-19 mice/group. $p$=0.0005 for D7, $p$=0.0162 for D21 by two-tailed t-test). (**C**) CD68 immunostain demonstrates that $\beta1^{AT2\text{-}KO}$ lungs contain increased macrophages 21 days post-LPS, quantified in (**D**), 20.6±2.9 CD68+ cells/ field in $\beta1^{AT2\text{-}KO}$ lungs versus 8.2±0.2 CD68+ cells/ field in $\beta1^{f/f}$ lungs (n=7 mice/ group, $p$=0.001, t=4.331, df-12). (**E**) Hematoxylin and eosinstained lung sections of $\beta1^{f/f}$ and $\beta1^{AT2\text{-}KO}$ mice administered doxycycline 5 days after intratracheal LPS dose, demonstrating emphysema at 21 days after LPS in the absence of acute inflammation in $\beta1^{AT2\text{-}KO}$ lungs. (**F**) Mean linear intercept quantified emphysematous alveolar remodeling in delayed doxycycline $\beta1^{AT2\text{-}KO}$ lungs at 21 days post-LPS, 28.3±1.3 μm in $\beta1^{f/f}$ lungs vs. 37.5±1.8 μm in $\beta1^{AT2\text{-}KO}$ lungs (n=6 mice/ group, $p$=0.0017 by two-tailed t-test). * $p < 0.05$. Scale bar = 100 μm for C and high-power insets in E; scale bar = 200 μm for low-power fields in E.

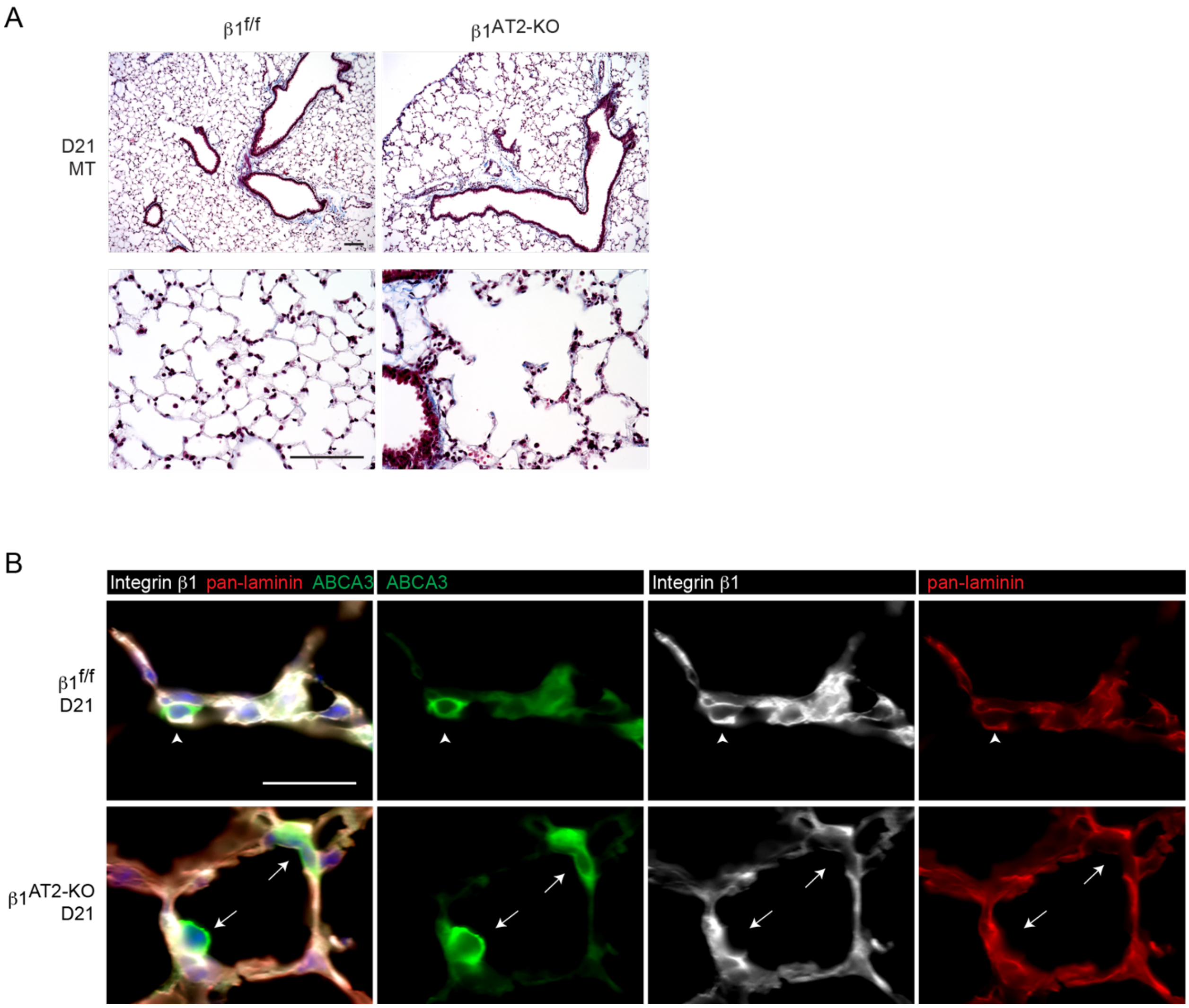


**Supplemental Figure 2. β1$^{AT2-KO}$ mice do not develop fibrosis from LPS-induced lung injury.** (**A**) Lung sections from β1$^{f/f}$ and β1$^{AT2-KO}$ mice with Masson's trichrome staining (MT) show no obvious fibrotic phenotype at 21 days after LPS treatment. (**B**) Lung sections immunostained for integrin β1 (white), pan-laminin (red), and the AT2 marker ABCA3 (green) demonstrate continued loss of β1 in AT2 cells 21 days after LPS treatment in β1$^{AT2-KO}$ lungs (arrows), while β1$^{f/f}$ AT2 cells express β1 (arrowheads). The basement membrane in β1$^{f/f}$ and β1$^{AT2-KO}$ lungs exhibit comparable pan-laminin content and localization 21 days after LPS treatment. Scale bar = 100 μm for A (both low and high-power images); scale bar = 25 μm for B.

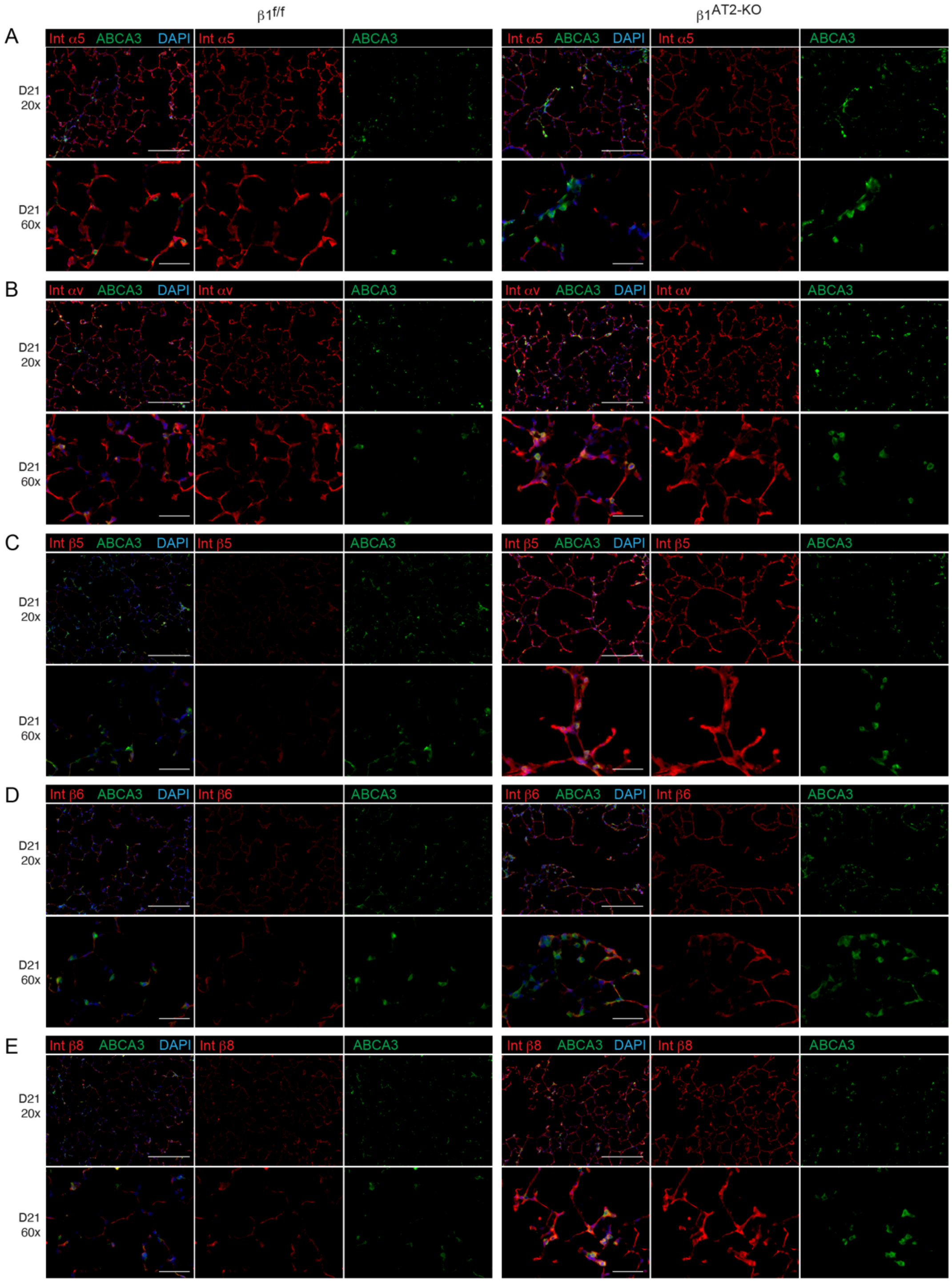


**Supplemental Figure 3. β1$^{AT2\text{-}KO}$ mice exhibit increased expression of non-β1-containing RGD-binding integrin subunits in late repair.** β1$^{f/f}$ and β1$^{AT2\text{-}KO}$ lung sections harvested 21 days after LPS were immunostained for integrin subunits (red, as above) and the AT2 marker ABCA3 (green). Scale bar = 200 μm for 20x images and 50 μm for 60x images.

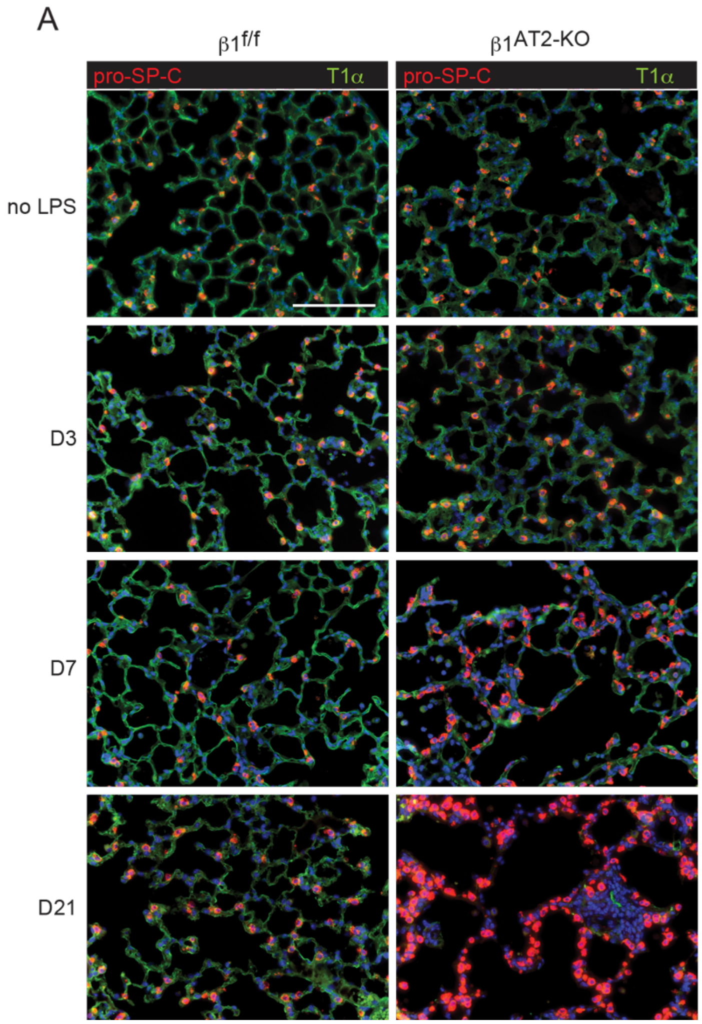


**Supplemental Figure 4. β1$^{AT2-KO}$ mice accumulate increased pro-SP-C+ AT2 cells and decreased AT1 marker T1α throughout repair.** (**A**) Lung sections immunostained for pro-SP-C (red) and T1α (green) demonstrate increased AT2 cells and decreased T1α in β1$^{AT2-KO}$ lungs by day 7 (D7) post-injury. At day 21 after LPS (D21), pro-SP-C+ cover the alveolar septa in heavily injured areas in β1$^{AT2-KO}$ lungs. Scale bar = 100 μm for A.

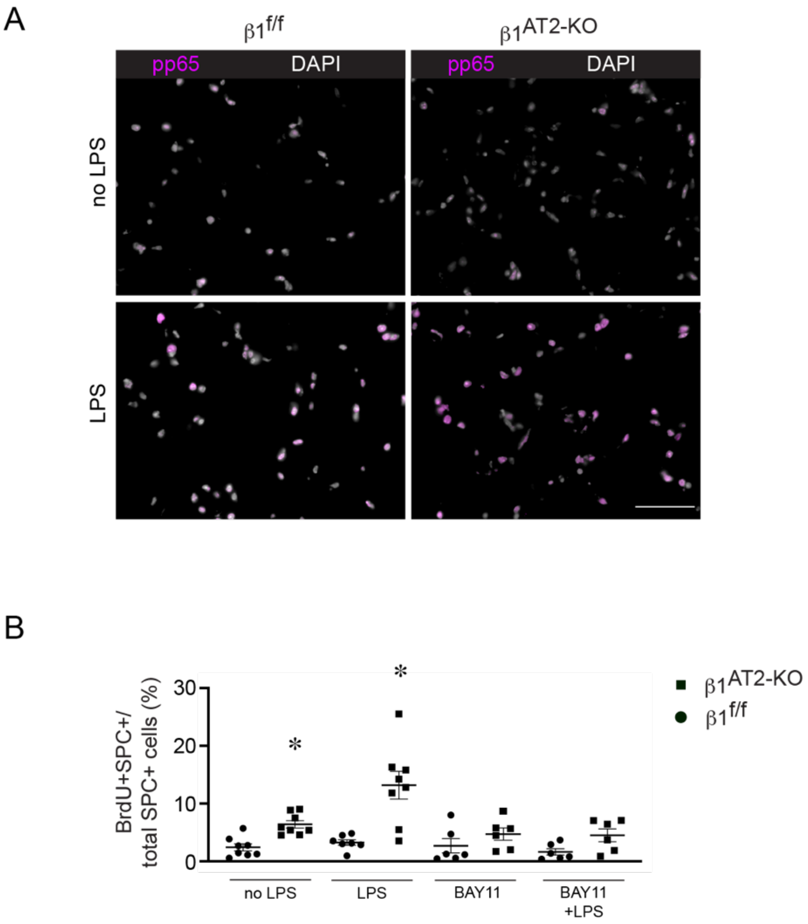


**Supplemental Figure 5. LPS-induced inflammation induced *ex vivo* in PCLS.** (A) PCLS treated with 62.5 ng/ml LPS induced NF-κB activation *ex vivo* as detected by nuclear phospho-p65 immunostain (pp65 in magenta, DAPI in white). (B) Individual values for proliferating (BrdU+) AT2 cells (pro-SP-C+) as a percent of total number of pro-SP-C+ cells. This data is presented as grouped line data with analysis by two-way ANOVA for genotype and treatment in Figure 3E, with statistics in Supplemental Table 1. * $p < 0.05$. Scale bar = 50 μm for A.

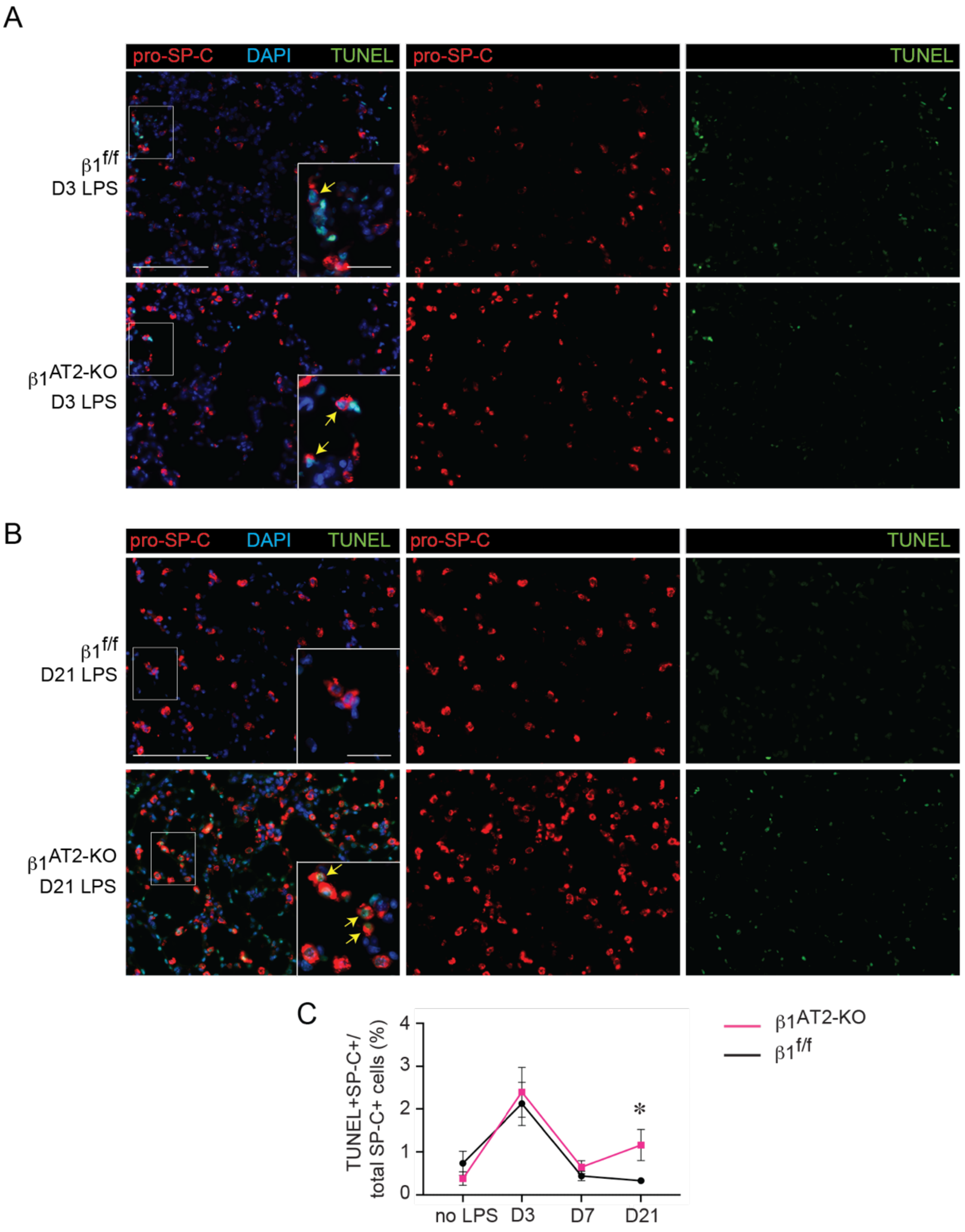


**Supplemental Figure 6. Increased apoptotic AT2 cells in β1$^{AT2\text{-}KO}$ mice in late repair.** (**A-B**) Immunodetection for apoptotic (by TUNEL assay) pro-SP-C+ AT2 cells at day three (D3) and day 21 (D21) after LPS treatment. (**C**) Quantification of TUNEL/pro-SP-C dual + cells per total number pro-SP-C+ cells shows a modest increase in apoptotic AT2 cells at D21 in β1$^{AT2\text{-}KO}$ lungs compared to β1$^{f/f}$ lungs (n=6-18 mice/ group, 10 sections/ mouse; *p*=0.312 for no LPS mice; *p*=0.7348 at D3; *p*=0.2748 at D7; *p*=0.0414 at D21). * $p < 0.05$. Scale bar = 100 μm for low power in A and B; scale bar = 25 μm for insets in A and B. Two-tailed t-test was used to compare genotypes at each time point in C.

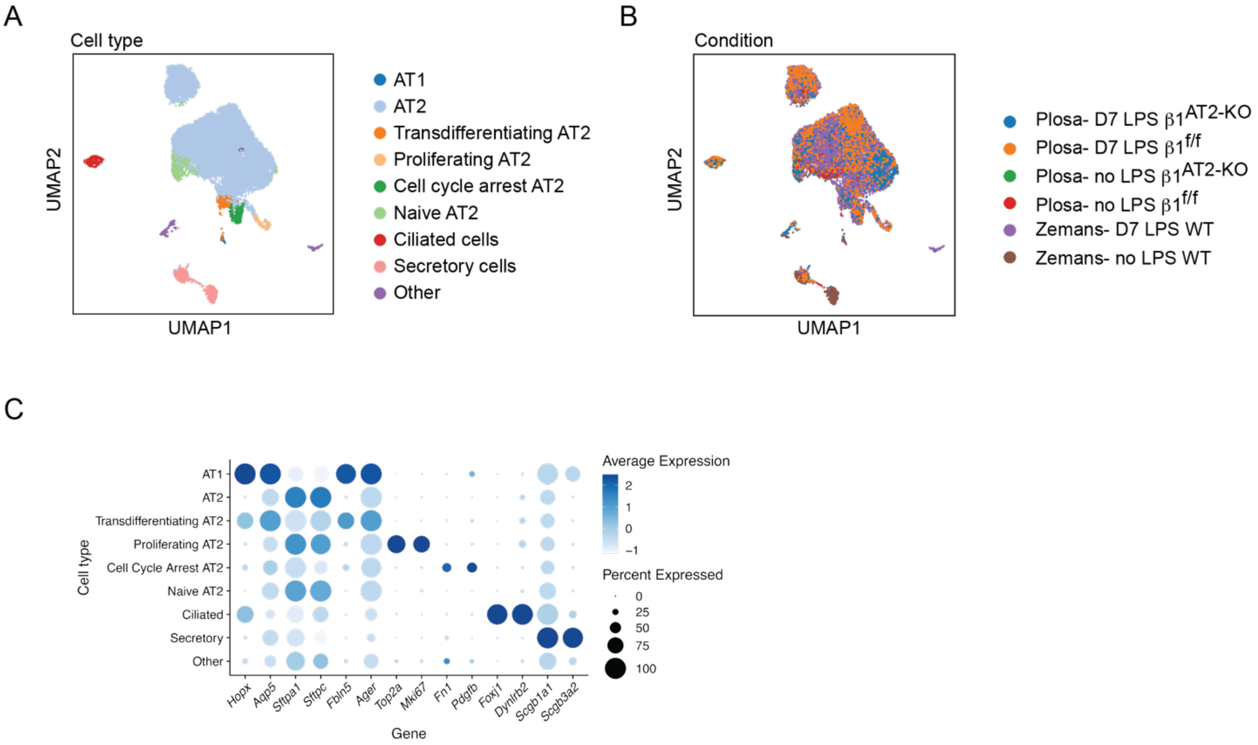


**Supplemental Figure 7**. **Integration of single-cell sequencing data from this study and from Riemondy et al. 2019 indicates that similar cell types are present in both studies**. Cell types were annotated according to marker gene expression patterns from Riemondy et al. 2019. (**A**) UMAP embedding of the integrated dataset with 15,154 cells is colored by cell type. (**B**) UMAP embedding colored by condition where cells from this study (Plosa), Riemondy et al. 2019 (Zemans), where each condition has individual colors (wild-type mice denoted WT). (**C**) Marker gene expression for each annotated cell type displayed in a dot plot where higher expression is represented as a darker color. The size of the dot indicates proportion of cells expressing each marker.

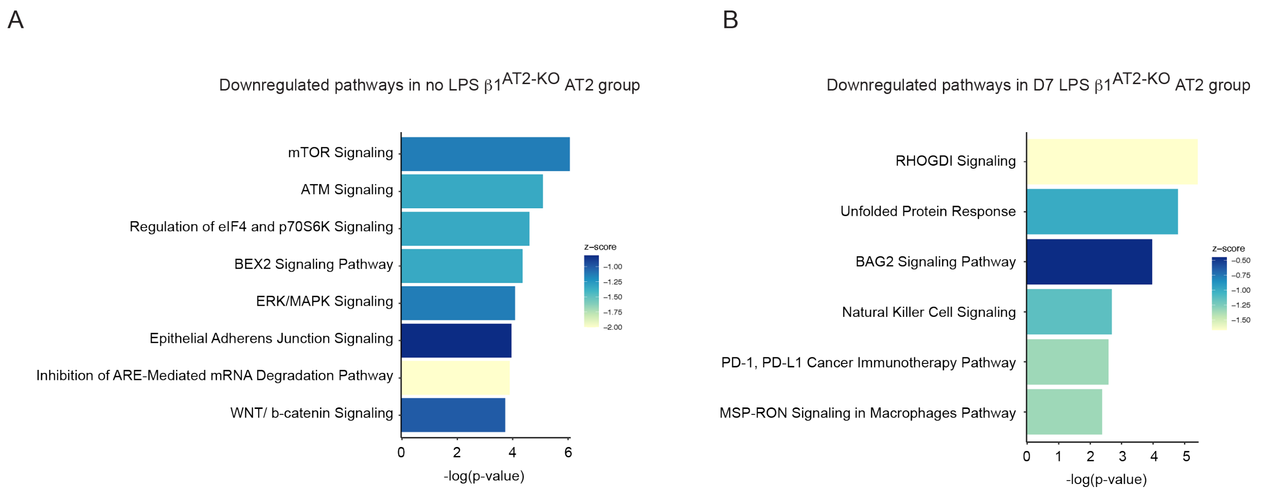


**Supplemental Figure 8. Downregulated pathways identified by Ingenuity Pathway Analysis.** (**A**) Downregulated pathways identified by Ingenuity Pathway Analysis on AT2 group from uninjured $\beta1^{f/f}$ and $\beta1^{AT2\text{-}KO}$ lungs. (**B**) Downregulated pathways by Ingenuity Pathway Analysis on AT2 group from $\beta1^{f/f}$ and $\beta1^{AT2\text{-}KO}$ lungs at day 7 after LPS injury.

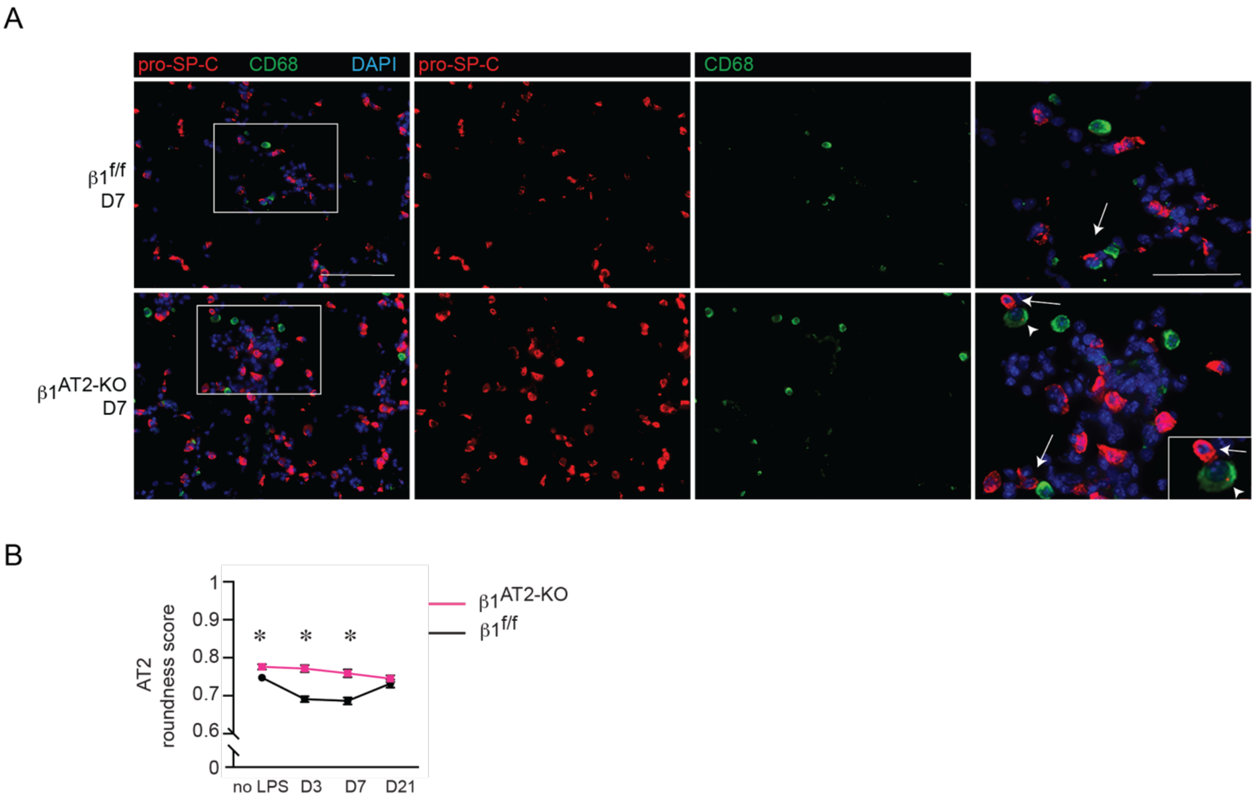


**Supplemental Figure 9. Roundness score is calculated from pro-SP-C+/ CD68- cells (AT2 cells exclusive of efferocytosing macrophages) at seven days after LPS treatment.** (**A**) Representative images of day seven (D7) $\beta1^{AT2\text{-}KO}$ and $\beta1^{f/f}$ lung sections immunostained for pro-SP-C (red) and CD68 (green) demonstrates few instances of macrophages efferocytosing AT2 cells (denoted by arrowhead) versus closely adjacent AT2 cells to macrophages (arrows). D7 cell shape metrics are based on this immunostaining (Figure 5C-D). (**B**) Roundness score calculated from pro-SP-C+ cells from uninjured, day 3, day 7, and day 21 LPS-treated $\beta1^{f/f}$ and $\beta1^{AT2\text{-}KO}$ mice (≥ 35 cells measured/ mouse from at least 3 different sections, n=6-8 mice/ group, two-tailed t-test comparing genotypes at each time point, $p$=0.0222 for uninjured mice; $p$<0.0001 at D3; $p$=0.0003 at D7; $p$=0.3641 at D21). ∗ $p < 0.05$. Scale bar = 100 μm for low-power in A, 50 μm for inset.

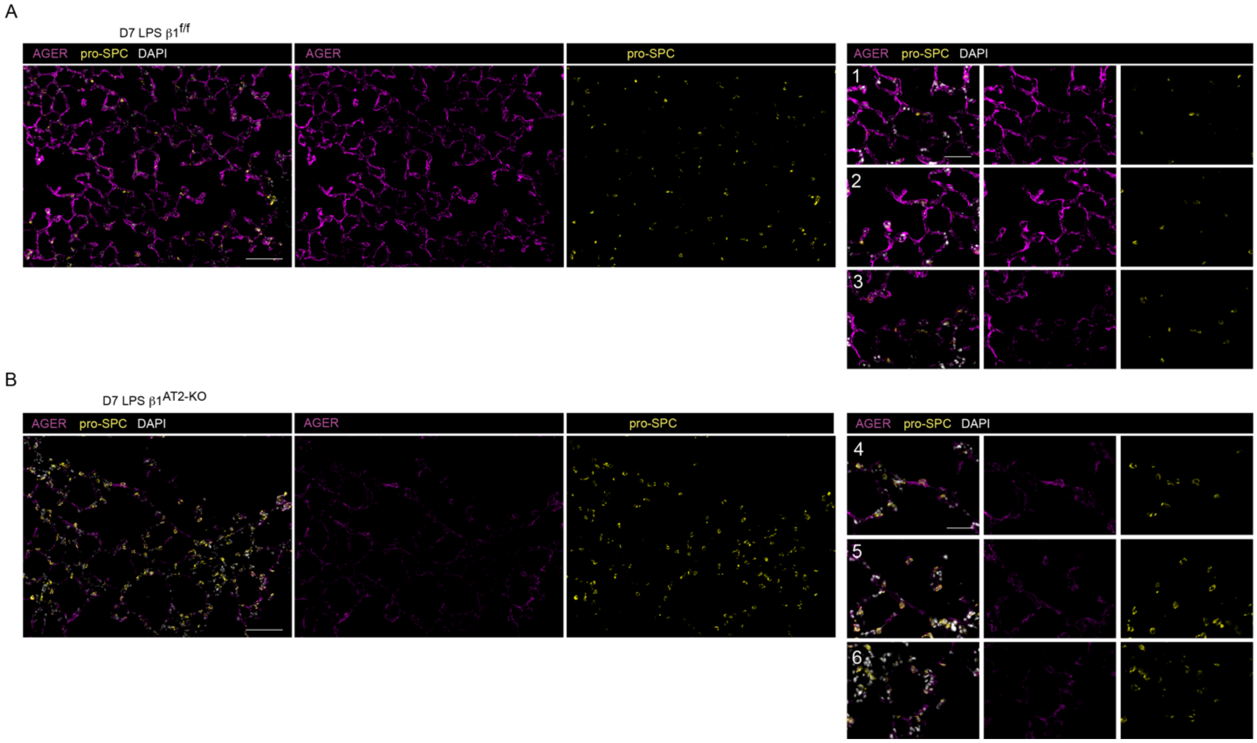


**Supplemental Figure 10.** (**A-B**) Representative single channel low-power images of day seven $\beta1^{f/f}$ (A) and $\beta1^{AT2\text{-}KO}$ (B) lung sections immunostained for AGER (purple) and pro-SP-C (gold) with DAPI nuclear probe (white). Scale bar = 100 µm for low-power in A and B, 50 µm for numbered inset.

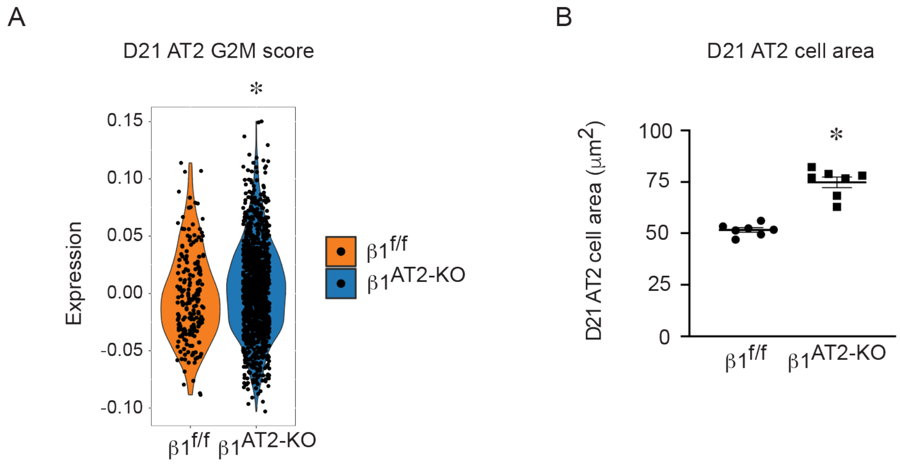


**Supplemental Figure 11.** (**A**) G2M proliferation score demonstrates sustained AT2 proliferation at 21 days after LPS injury in $\beta1^{AT2\text{-}KO}$ lungs (mean G2M score 0.00385793±0.03973903 for $\beta1^{AT2\text{-}KO}$ AT2 cells vs. -0.0019844±0.036727215 for $\beta1^{f/f}$ AT2 cells, # $p = 0.01789$). (**B**) Area of pro-SP-C+/CD68- AT2 cells from D21 LPS-treated $\beta1^{f/f}$ and $\beta1^{AT2\text{-}KO}$ mice (51.5±1.1 $\mu m^2$ in $\beta1^{f/f}$ lungs compared to 74.9±2.6 $\mu m^2$ in $\beta1^{AT2\text{-}KO}$ lungs, n = 7 $\beta1^{f/f}$ and 7 $\beta1^{AT2\text{-}KO}$ mice, ≥ 50 cells measured/ mouse; cells analyzed from 5 different sections, two-tailed t-test, $p$=0.0001).

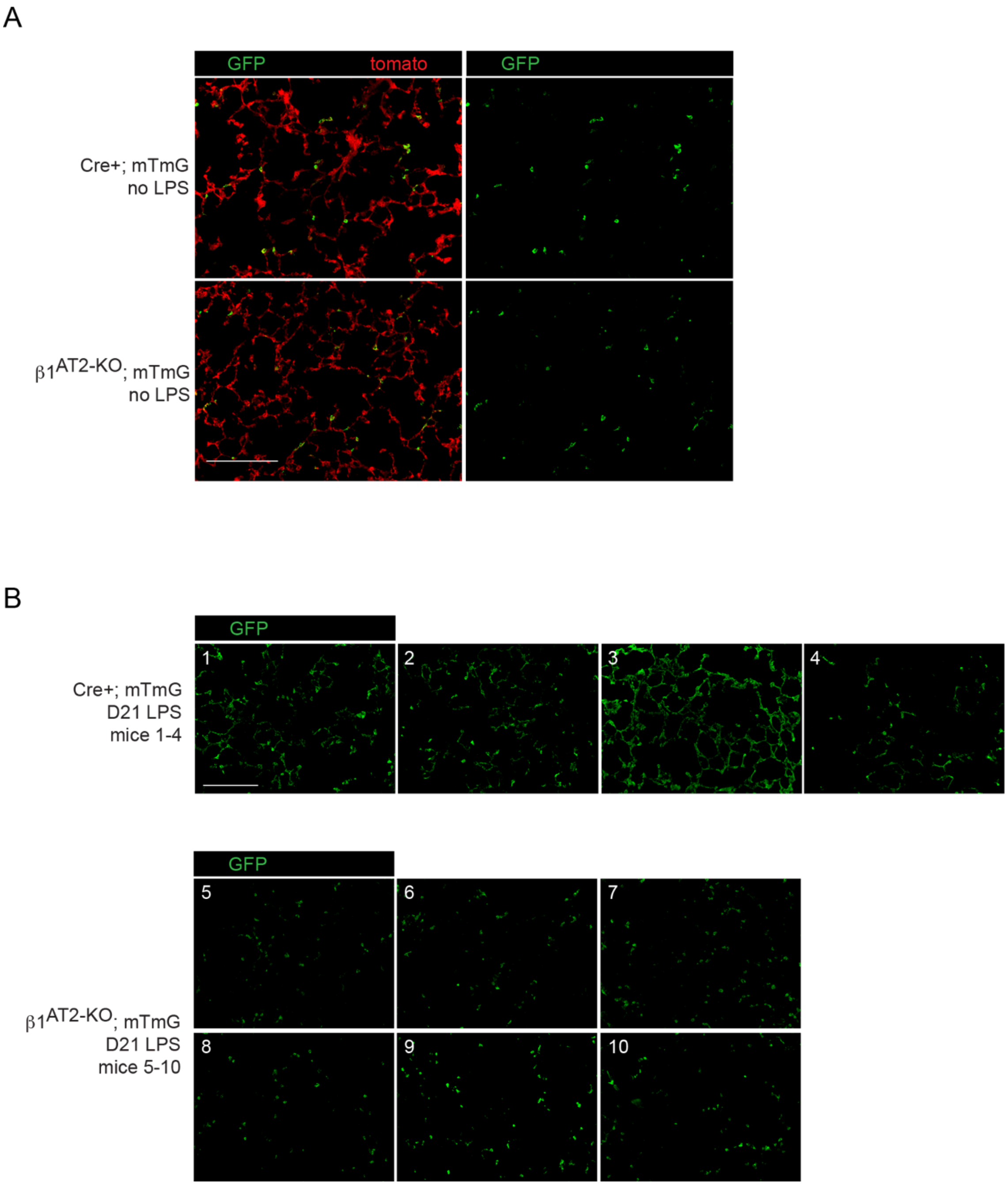


**Supplemental Figure 12.** (**A**) Merge and single channel GFP panel images of uninjured Cre+; mTmG and $\beta1^{AT2\text{-}KO}$; mTmG mice. Images demonstrate that mTmG Cre-recombinase reporter labels cells of AT2 morphology only prior to injury in both control and $\beta1^{AT2\text{-}KO}$; mTmG mice. (**B**) Representative images of GFP channel from mTmG labeled mice demonstrate GFP label on cells of both AT2 and AT1 morphology in control mTmG mice 21 days after LPS. However, $\beta1^{AT2\text{-}KO}$; mTmG mice uniformly possess only GFP labeled cells of round, AT2 morphology. Images from all mice are presented (control mice #1-4 and $\beta1^{AT2\text{-}KO}$; mTmG mice #5-10). Scale bar = 200 μm in A and B.

**Supplemental Table 1.**

Statistical analysis.

Two-tailed t-test

| Figure | Timepoint | Sample size | | $p$-value | t-values | df |
|---|---|---|---|---|---|---|
| 1B | D21 | 7 $\beta1^{f/f}$ | 6 $\beta1^{AT2\text{-}KO}$ | 0.0014 | 4.229 | 11 |
| 1C | No LPS | 7 $\beta1^{f/f}$ | 6 $\beta1^{AT2\text{-}KO}$ | 0.0485 | 2.218 | 11 |
| | D3 | 12 $\beta1^{f/f}$ | 7 $\beta1^{AT2\text{-}KO}$ | 0.0036 | 3.375 | 17 |
| | D7 | 11 $\beta1^{f/f}$ | 12 $\beta1^{AT2\text{-}KO}$ | 0.0050 | 3.132 | 21 |
| | D21 | 14 $\beta1^{f/f}$ | 9 $\beta1^{AT2\text{-}KO}$ | 0.2628 | 1.151 | 21 |
| 1D | No LPS | 7 $\beta1^{f/f}$ | 6 $\beta1^{AT2\text{-}KO}$ | 0.0002 | 5.541 | 11 |
| | D3 | 10 $\beta1^{f/f}$ | 7 $\beta1^{AT2\text{-}KO}$ | 0.0730 | 1.928 | 15 |
| | D7 | 26 $\beta1^{f/f}$ | 22 $\beta1^{AT2\text{-}KO}$ | 0.0007 | 3.622 | 46 |
| | D21 | 14 $\beta1^{f/f}$ | 8 $\beta1^{AT2\text{-}KO}$ | <0.0001 | 5.686 | 20 |
| 2C | D21 | 8 $\beta1^{f/f}$ | 8 $\beta1^{AT2\text{-}KO}$ | 0.0086 | 3.055 | 14 |
| 2D | D21 | 6 $\beta1^{f/f}$ | 6 $\beta1^{AT2\text{-}KO}$ | 0.0003 | 5.487 | 10 |
| 3A | No LPS | 7 $\beta1^{f/f}$ | 6 $\beta1^{AT2\text{-}KO}$ | 0.0247 | 2.599 | 11 |
| | D3 | 6 $\beta1^{f/f}$ | 6 $\beta1^{AT2\text{-}KO}$ | 0.8220 | 0.2304 | 11 |
| | D7 | 7 $\beta1^{f/f}$ | 8 $\beta1^{AT2\text{-}KO}$ | 0.0001 | 5.456 | 13 |
| | D21 | 8 $\beta1^{f/f}$ | 8 $\beta1^{AT2\text{-}KO}$ | 0.0009 | 4.190 | 14 |
| 3B | No LPS | 7 $\beta1^{f/f}$ | 6 $\beta1^{AT2\text{-}KO}$ | 0.0003 | 5.212 | 11 |
| | D3 | 7 $\beta1^{f/f}$ | 8 $\beta1^{AT2\text{-}KO}$ | 0.0311 | 2.417 | 13 |
| | D7 | 7 $\beta1^{f/f}$ | 8 $\beta1^{AT2\text{-}KO}$ | 0.0310 | 2.419 | 13 |
| | D21 | 8 $\beta1^{f/f}$ | 8 $\beta1^{AT2\text{-}KO}$ | 0.0128 | 2.853 | 14 |
| 5C | D7 | 6 $\beta1^{f/f}$ | 6 $\beta1^{AT2\text{-}KO}$ | <0.0001 | 6.715 | 10 |
| 5D | D7 | 6 $\beta1^{f/f}$ | 6 $\beta1^{AT2\text{-}KO}$ | 0.0009 | 4.658 | 10 |
| 5F | D7-JLA20 | 6 $\beta1^{f/f}$ | 5 $\beta1^{AT2\text{-}KO}$ | 0.0088 | 3.329 | 9 |
| 5G | D7-phalloidin | 6 $\beta1^{f/f}$ | 5 $\beta1^{AT2\text{-}KO}$ | 0.0482 | 2.285 | 9 |
| 7F | D21 | 6 $\beta1^{f/f}$ | 6 $\beta1^{AT2\text{-}KO}$ | <0.0001 | 6.638 | 10 |
| 8B | D21 | 4 Cre+; mTmG | 6 $\beta1^{AT2\text{-}KO}$; mTmG | 0.0087 | 3.448 | 8 |

One-way ANOVA

| Figure | Sample size | | | | F value | df |
|---|---|---|---|---|---|---|
| 5I | 6 $\beta1^{f/f}$ no LPS | 7 $\beta1^{AT2\text{-}KO}$ no LPS | 8 $\beta1^{f/f}$ D7 | 7 $\beta1^{AT2\text{-}KO}$ D7 | 53.42 | 3 |
| 5J | 6 $\beta1^{f/f}$ no LPS | 7 $\beta1^{AT2\text{-}KO}$ no LPS | 8 $\beta1^{f/f}$ D7 | 7 $\beta1^{AT2\text{-}KO}$ D7 | 17.46 | 3 |
| 5K | 6 $\beta1^{f/f}$ no LPS | 7 $\beta1^{AT2\text{-}KO}$ no LPS | 8 $\beta1^{f/f}$ D7 | 7 $\beta1^{AT2\text{-}KO}$ D7 | 31.09 | 3 |

Two-way ANOVA

| Figure | Sample size | No LPS | LPS | BAY 11 | Bay 11+LPS | | $p$-value | F value | df |
|---|---|---|---|---|---|---|---|---|---|
| 3E | $\beta1^{f/f}$ | 8 | 7 | 6 | 6 | treatment | 0.0010 | 6.3 | 3 |
| | $\beta1^{AT2\text{-}KO}$ | 8 | 8 | 6 | 6 | genotype | <0.0001 | 26.1 | 1 |
